# Modular basis for potent SARS-CoV-2 neutralization by a prevalent VH1-2-derived antibody class

**DOI:** 10.1101/2021.01.11.426218

**Authors:** Micah Rapp, Yicheng Guo, Eswar R. Reddem, Lihong Liu, Pengfei Wang, Jian Yu, Gabriele Cerutti, Jude Bimela, Fabiana Bahna, Seetha Mannepalli, Baoshan Zhang, Peter D. Kwong, David D. Ho, Lawrence Shapiro, Zizhang Sheng

## Abstract

Antibodies with heavy chains that derive from the VH1-2 gene constitute some of the most potent SARS-CoV-2-neutralizing antibodies yet identified. To provide insight into whether these genetic similarities inform common modes of recognition, we determined structures of the SARS-CoV-2 spike in complex with three VH1-2-derived antibodies: 2-15, 2-43, and H4. All three utilized VH1-2-encoded motifs to recognize the receptor-binding domain (RBD), with heavy chain N53I enhancing binding and light chain tyrosines recognizing F486_RBD_. Despite these similarities, class members bound both RBD-up and -down conformations of the spike, with a subset of antibodies utilizing elongated CDRH3s to recognize glycan *N*343 on a neighboring RBD – a quaternary interaction accommodated by an increase in RBD separation of up to 12 Å. The VH1-2-antibody class thus utilizes modular recognition encoded by modular genetic elements to effect potent neutralization, with VH-gene component specifying recognition of RBD and CDRH3 component specifying quaternary interactions.

**Highlights:**

- Determine structures of VH1-2-derived antibodies 2-43, 2-15, and H4 in complex with SARS-CoV-2 spike
- Define a multi-donor VH1-2-antibody class with modular components for RBD and quaternary recognition
- Reveal structural basis of RBD-up and RBD-down recognition within the class
- Show somatic hypermutations and avidity to be critical for potency
- Delineate changes in spike conformation induced by CDRH3-mediated quaternary recognition

## Introduction

Studies on human antibody responses to viral pathogens including HIV-1, influenza, EBOLA, and malaria have revealed prominent classes of similar neutralizing antibodies (nAbs), which arise commonly in numerous individuals in response to infection or vaccination (Zhou et al., 2013, Ehrhardt et al., 2019, Zhou et al., 2015, Joyce et al., 2016, Imkeller et al., 2018, Kallewaard et al., 2016, Pappas et al., 2014). Such multi-donor antibody classes are defined based on similar V(D)J gene recombination and similar mode of structural recognition – with this combination indicative of a common evolutionary process in antibody development (Kwong and Mascola, 2012). Multi-donor antibody classes are thought to arise based on effective function, combined with genetic accessibility due to class requirements for V(D)J recombination and somatic hypermutations of sufficient frequency as to be present in the antibody human repertoire. Some multi-donor antibody classes are potent, broadly neutralizing, and frequent in human antibody repertoire (Joyce et al., 2016, Zhou et al., 2013, Imkeller et al., 2018). A prominent mode of rational vaccine design – “lineage-based vaccine design” –endeavors to elicit such antibody classes by vaccination (Kwong and Mascola, 2018, Jardine et al., 2013, Haynes et al., 2012) and this approach to human vaccination has recently entered clinical assessment (Diemert and McElrath, 2018).

Severe Acute Respiratory Syndrome Coronavirus 2 (SARS-CoV-2), the causative agent of the ongoing Coronavirus disease 2019 (COVID-19) pandemic, has infected over 80 million people and has claimed over one million deaths since the outbreak began in late 2019 (Zhu et al., 2020, Zhou et al., 2020, Dong et al., 2020). Therapies and vaccines are urgently needed to end the pandemic. Many nAbs have now been isolated from COVID-19 convalescent donors with the most potent nAbs showing promise as prophylactic or therapeutic agents (Liu et al., 2020, Robbiani et al., 2020, Rogers et al., 2020, Brouwer et al., 2020, Zost et al., 2020, Kreer et al., 2020, Tortorici et al., 2020, Chen et al., 2020). This growing set of nAbs provides an opportunity to identify effective human antibody responses to SARS-CoV-2 common in the population, which will inform therapeutic strategies and help to interpret vaccine readouts.

SARS-CoV-2 nAbs predominantly target the viral spike glycoprotein, which interacts with angiotensin-converting enzyme 2 (ACE2) receptors on the host-cell surface to mediate virus entry (Hoffmann et al., 2020, Wang et al., 2020). The ectodomain of the prefusion spike comprises three copies of both S1 and S2 subunits (Wrapp et al., 2020). The S1 subunit is responsible for ACE2 binding, and the S2 subunit mediates fusion with host cell membrane. Each S1 subunit comprises an N-terminal domain (NTD) and a receptor binding domain (RBD). The RBDs are very flexible, adopting either an ‘up’ conformation (open state) or a ‘down’ conformation (closed state), with only the ‘up’ RBDs capable of interacting with ACE2 (Wang et al., 2020). Currently, many nAbs have been characterized which bind to epitopes on either the ‘up’ and/or ‘down’ RBDs (Tortorici et al., 2020, Liu et al., 2020, Barnes et al., 2020a). These RBD-targeting nAbs neutralize SARS-CoV-2 through mechanisms that include competition with ACE2 for RBD binding and locking the RBDs in the ‘down’ conformation.

Human SARS-CoV-2 nAbs develop with few somatic hypermutations and strong avidity effects (Liu et al., 2020, Robbiani et al., 2020, Barnes et al., 2020b, Kreer et al., 2020). Characterization of genetic features of SARS-CoV-2 nAbs identified to date show enrichment of antibody variable genes including VH3-53, VH1-2, VH1-69, VH3-66, VH1-58, and VH3-30 (Yuan et al., 2020, Robbiani et al., 2020, Liu et al., 2020). So far, structural characterization of multiple nAbs have revealed two separate RBD-targeted classes derived from the similar VH3-53 and VH3-66 genes. One of these is characterized by heavy chain complementarity-determining region 3 (CDRH3) of 15 amino acids or shorter – and recognizes RBD ridge in the ‘up’ position on SARS-CoV-2 spike (type I) (Yuan et al., 2020). The second VH3-53/-66 class – with longer CDRH3s – recognizes a similar region of RBD, but adopts an approach angle with the heavy and light chain orientation rotated 180 degrees (type II), suggesting CDRH3 to be critical for the classification of VH3-53/-66 antibodies (Wu et al., 2020a, Barnes et al., 2020a). Different VH3-30 originated antibodies can recognize at least three different regions of RBDs (Barnes et al., 2020a, Hansen et al., 2020), demonstrating that they do not form a single gene-restricted antibody class. Nevertheless, whether nAbs derived from VH1-2 and other germline genes form gene-restricted classes that represent shared effective antibody responses remains unaddressed. Currently, three VH1-2 potent nAbs (2-4, S2M11, and C121) show similar modes of RBD recognition but differences in quaternary epitope recognition (Tortorici et al., 2020, Liu et al., 2020, Barnes et al., 2020a). It is thus still unclear whether RBD-targeting VH1-2 antibodies form a gene-restricted class. If so, what are the key genetic and structural signatures and determinants of neutralization potency?

Here we present structures of three VH1-2-derived nAbs (2-15, 2-43, and H4), revealing that they recognize a SARS-CoV-2 RBD epitope with similar modes of RBD recognition and similar angles of approach. Overall, recognition is modular with RBD predominantly recognized by a VH1-2 gene encoded module. The second recognition module is represented by the diverse CDRH3s of the VH1-2 antibodies, which mediate quaternary recognition of N343 glycan from an adjacent RBD for a subset of class members. Thus, we define a multi-donor VH1-2 antibody class, members of which can achieve very high neutralization potency, which is prevalent in human responses to SARS-CoV-2. The shared genetic and structural signatures inform strategies to improve members of the VH1-2 antibody class.

## Results

### VH1-2 antibodies are prevalent in human response to SARS-CoV-2 infection

To identify common features of the human antibody response to SARS-CoV-2, 158 spike-specific antibodies with characterized neutralization potencies were collected from 10 studies (Table S1) (Liu et al., 2020, Wu et al., 2020b, Hansen et al., 2020, Robbiani et al., 2020, Rogers et al., 2020, Brouwer et al., 2020, Ju et al., 2020, Zost et al., 2020, Kreer et al., 2020, Tortorici et al., 2020). V(D)J gene usage analysis showed that VH1-2 was the second most frequently utilized germline gene (Figure 1A, 25 in total). Comparison of neutralization potencies revealed that 11 of 25 (44%) of the VH1-2 antibodies are potent (half maximal inhibitory concentration (IC_50_) <0.1µg/ml, Figure 1B). Within four studies (Zost et al., 2020, Tortorici et al., 2020, Liu et al., 2020, Hansen et al., 2020), VH1-2 antibodies ranked the most potent of gene-delimited sets of antibodies (IC_50_ ranges from 0.015 to 0.0007µg/ml). All 25 of the VH1-2-derived antibodies have been reported to bind to the SARS-CoV-2 RBD (Liu et al., 2020, Wu et al., 2020b, Hansen et al., 2020, Robbiani et al., 2020, Rogers et al., 2020, Brouwer et al., 2020, Ju et al., 2020, Zost et al., 2020, Kreer et al., 2020, Tortorici et al., 2020). Sequence alignment of the VH1-2 nAbs showed the heavy chains to carry few somatic hypermutations, with each having a unique CDRH3 with length varying from 11 to 21 amino acids (Figure 1C and S1A, Kabat definition), and using different D genes (Figure S1B). The VH1-2 antibodies used both kappa and lambda genes with enrichment of the IGLV2-14 gene (Figure 1C).

**Figure 1.**
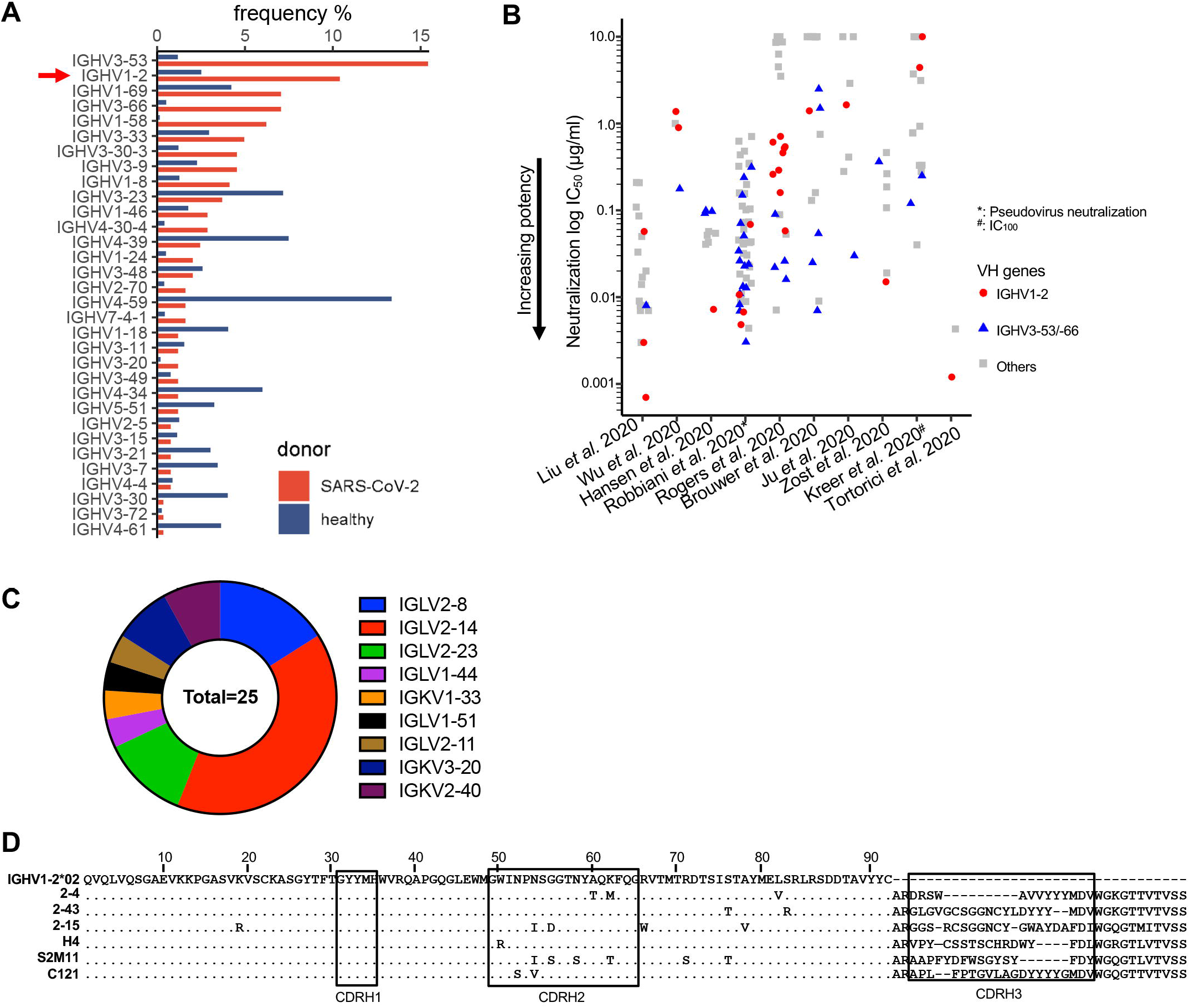
SARS-CoV-2 infection in humans induces potently neutralizing VH1-2 antibodies. (A) Gene usage of SARS-CoV-2 neutralizing antibodies. SARS-CoV-2 spike-specific VH1-2 antibodies are frequently induced in infected humans. VH1-2 antibodies are significantly enriched in the antigen-specific antibody repertoire than that of heathy donors. (B) Many SARS-CoV-2 neutralizing VH1-2 antibodies (red) isolated from human donors achieve high potency, comparable to the most frequent IGHV3-53/-66 antibodies (blue). The half maximal inhibitory concentration (IC_50_) of neutralization are shown except IC_100_ for antibodies from Kreer *et al*., (2020). Live virus neutralization potency is shown for antibodies from nine studies except the Hansen study. IC_50_ values greater than 10µg/ml are set to 10µg/ml. (C) SARS-CoV-2 neutralizing VH1-2 antibodies use diverse light chain genes. (D) Sequence alignment of the heavy chain of six VH1-2 antibodies that are characterized in this study. Residues identical to germline gene are shown as dots. See also Figure S1 and Table S1.

As described below, we determined structures of three nAbs: 2-15, 2-43, and H4, with the highly potent antibodies 2-15 and 2-43 isolated from donor ‘2’ of our previous study (Liu et al., 2020) while H4 was from a different donor (Wu et al., 2020b). 2-15, 2-43, and H4 neutralize authentic SARS-CoV-2 “live” virus with IC_50_ of 0.0007, 0.003, and 0.896 µg/ml, respectively. These three antibodies derived from different heavy and light chain gene recombinations, and hence three different B cell lineages (Figure S1B). Both 2-43 and 2-15 utilize the DH2-15*01 gene and have a long CDRH3 of 20 amino acids (Figure S1C), but they have different HJ gene origins (JH6*03 and JH3*02 respectively). The light chains of 2-43 and 2-15 were derived from recombination of a novel allele of the VL2-14 gene with JL3*02 and JL1*01 respectively (Figure S1D). H4 used DH2-2*01 and JH2*01 genes and had a CDRH3 of 17 amino acids (Wu et al., 2020b). The H4 light chain was derived from VK2-40*01 and JK4*01.

### Structures of antibodies 2-15, 2-43, and H4 in complex with SARS-CoV-2 spike or RBD

To understand SARS-CoV-2 spike recognition by the VH1-2-derived antibodies, we used single-particle cryo-electron microscopy (cryo-EM) to produce high-resolution 3D-reconstructions of antigen-binding fragments (Fabs) from 2-15, 2-43, and H4 in complex with the SARS-CoV-2 spike (Table S2). The reconstruction of the 2-43 complex with spike, refined to a global resolution of 3.60Å (Figure S2A-S2E), is significantly improved than in our previous study (5.8Å resolution) (Liu et al., 2020). The reconstruction revealed a predominant class with 3 Fab molecules bound per spike trimer (Figure 2A). Each 2-43 Fab used heavy chain variable domain to bind one primary RBD with an orientation similar to the previously published antibody 2-4 (Liu et al., 2020), with all RBDs in the ‘down’ conformation (Figure 2A). 2-43 heavy and light chains also recognize a quaternary epitope from the adjacent RBD. 3D classification revealed less populated states with 1 and 2 Fabs bound, but in every case the Fab was bound to a ‘down’ RBD. This dependence upon the ‘down’ conformation is likely due to extensive interactions between the Fab CDRH3 loop and the N-linked glycosylation at residue N343 on the adjacent RBD (Figure S2E). This suggests a neutralization mechanism whereby the Fab simultaneously occludes ACE2 binding and locks the RBDs in the ‘down’ conformation.

**Figure 2.**
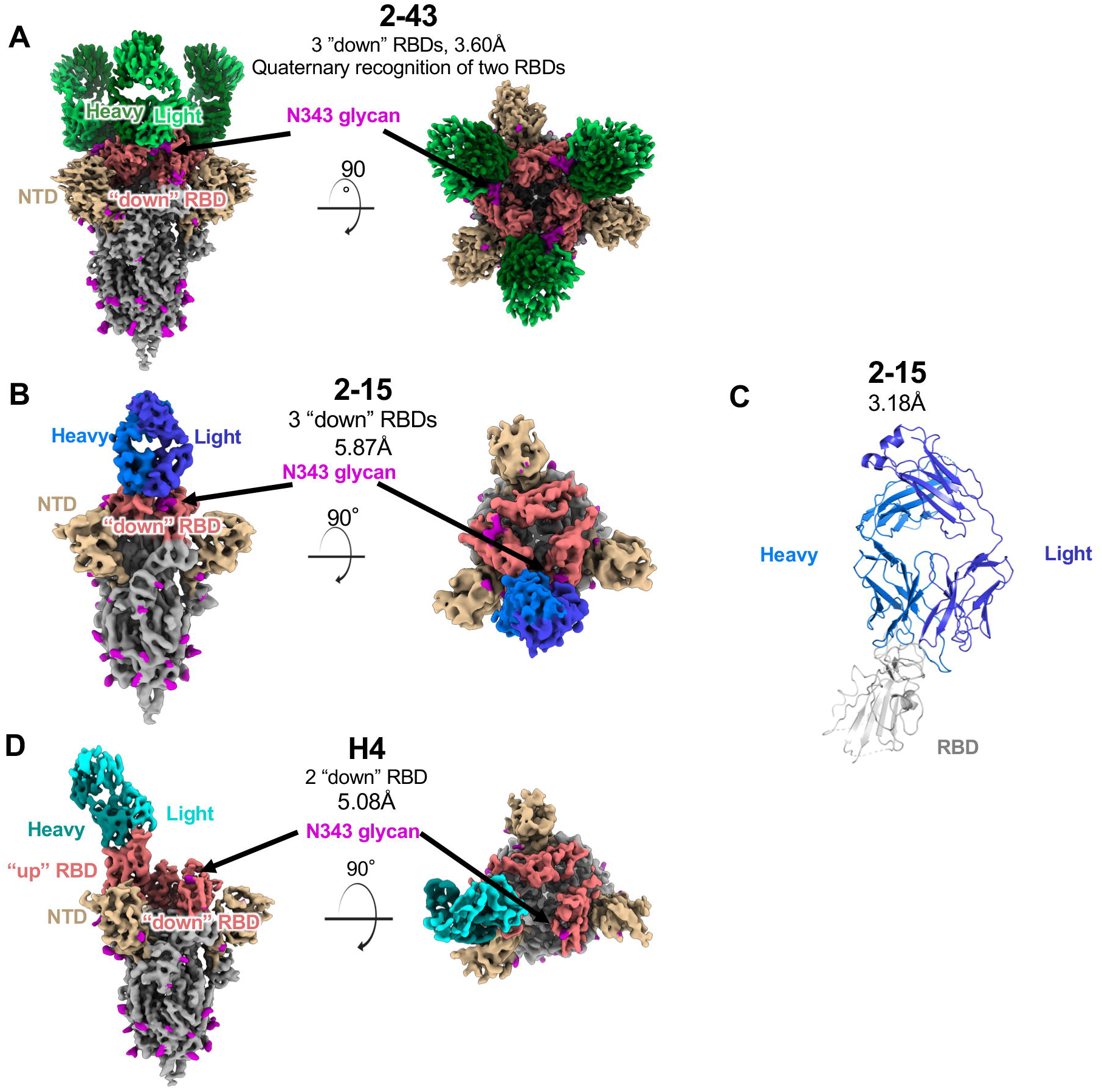
Structures of three SARS-CoV-2 neutralizing VH1-2 antibodies reveal both ‘RBD-down’ and ‘RBD-up’ modes of spike recognition. (A) Side and top views of three 2-43 antibody Fabs bound to prefusion SARS-COV-2 spike in the closed state. Color schemes are: 2-43 green; RBD, salmon; NTD, yellow; N-linked glycans, magenta; other spike regions, gray. (B) Side and top views of one 2-15 antibody Fabin complex with prefusion SARS-COV-2 spike in the closed state. 2-15 is colored blue. (C) Overview of the crystal structure of 2-15 in complex with SARS-CoV-2 RBD. Residue segments missing in the structure are shown as dashed lines. (D) Side and top views of one H4 Fab recognizing an “up” RBD on prefusion SARS-COV-2 spike. H4 is colored cyan. See also Figure S2, Table S2, and Table S3.

For the complex of 2-15 with spike (Figure 2B), approximately 56% of particles were bound to an RBD in the ‘up’ conformation, and 44% bound to RBD ‘down’ (Figure S2F-S2I), differing from antibody 2-43, which bound only to “down” RBDs. Due to increased conformational heterogeneity of the RBD ‘up’ class, the RBD ‘down’ class was the focus of our structural analysis (Figure 2B). In our initial attempt, a 9-fold molar excess of Fab was incubated with the S trimer. However, this resulted in spike disassembly (Figure S2J). To overcome this issue for structure determination, we found that a 1:1 molar ratio left spike-complexes intact, though some spike disassembly was still observed (Figure S2K). Although we were unable to resolve the 2-15:spike interaction at atomic resolution, the cryo-EM analysis did reveal the overall orientation of the Fab along with the position of the peptide backbone for several of the CDR loops.

To understand the binding mode of 2-15 at atomic resolution, we determined the crystal structure of 2-15 in complex with isolated RBD. The structure was determined by molecular replacement at 3.18 Å resolution to a final crystallographic R_work_/R_free_ of 18.6%/23.8% with good overall geometry (**Table S3)**. The electron density for RBD (corresponding to S1-subunit residues 333-527) was well defined for residues 344-518, with missing internal stretches of four (362-366) and seven (383-390) residues, with the C-terminal eighteen residues (519-537) disordered (Figure 2C). For Fab 2-15 all residues were visualized in density, with the exception of heavy chain residues 142-145 in the Fc region. We then docked the crystal structure to the 2-15-spike complex cryo-EM density map by superposing RBD from the crystal structure on RBD from the cryo-EM map, with subsequent rigid body fitting to density.

Overall, the conformation of 2-15 in the crystal structure agrees with the density observed in the cryo-EM map of the complex with spike – particularly in the V-gene-encoded CDR regions. However, the conformation of the long CDRH3 loop differs, and is far more elongated in the crystal structure (Figure S2L). In the cryo-EM map, it appears that the CDRH3 loop is pushed away from the N343 glycan and the helix encompassing residues 364-371 from the adjacent ‘down’ RBD. The two light chains exhibit more significant conformational differences. Save for CDRL3, which bends slightly away from RBD in the crystal structure, the conformations of the loops are quite similar, but the chain is shifted down closer to the RBD.

Finally, a reconstruction of Fab H4 revealed spike complexes with a single Fab bound to RBD in the ‘up’ conformation (Figure 2D and S2M to S2P). Interestingly, no Fab was seen bound to a ‘down’ RBD. The conformational flexibility of the RBD while in the ‘up’ conformation made a high-resolution cryo-EM reconstruction unattainable, and the map was refined to an overall resolution of 5.08Å. Similar to Fab 2-15, the peptide backbone for many CDR loops are observed, and a homology model of the Fab variable domain was docked into the density. The superimposition of modeled H4 to the 2-43/spike complex showed that H4 adopts an RBD approaching angle similar to 2-43 (Figure S2Q). However, the light chain of H4 rotates toward the interface between RBDs such that the long CDRL1 of H4 clashes with N343 glycan from an adjacent ‘down’ RBD (Figure S2R), which suggests a possible explanation for the apparent lack of H4 binding to ‘down’ RBDs.

### Conserved mode of RBD recognition

To understand the similarity in RBD recognition, we characterized the epitope and paratope interactions of 2-43 and 2-15 and compared to the three published VH1-2 antibodies 2-4, S2M11, and C121 (Tortorici et al., 2020, Liu et al., 2020, Barnes et al., 2020a). Overall, 2-43 interacts predominantly with the receptor binding motif (RBM, residues 438-508) on one ‘down’ RBD protomer while simultaneously binding the N343 glycan from an adjacent ‘down’ RBD protomer (Figure 3A). We describe the N343 glycan interaction below in the last section focused on quaternary interactions. The interaction between 2-43 and the ‘primary’ RBD bound through its receptor binding motif buries a total of 756Å^2^ paratope surface area (BSA), 83% of which is contributed by heavy chain (Figure 3A and S3A). Heavy chain framework 1 (FWH1), CDRH1, CDRH2, and CDRL3 of 2-43 form hydrogen bond networks with three RBD regions. In the first region, residues in FWH1 (G26_HC_, Y27_HC_, T28_HC_, F29_HC_, and T30_HC_) and the DE-loop in FWH3 (T73_HC_, S74_HC_, and T76_HC_) form a groove to hold Y449_RBD_ which contributes 111Å^2^ BSA. Hydrogen bonds are observed between the backbone atoms of Y27_HC_ and F29_HC_ and the side chain hydroxyl of Y449_RBD_ and between the side chain hydroxyl of T28_HC_ and the side chain amino group of Q498_RBD_ (Figure 3A). In the second region, involving the ‘flat’ region of RBM, the hydroxyl group of T30_HC_ forms a hydrogen bond with the backbone carbonyl from S494_RBD_. In the third region or the RBD ridge region, Y33_HC_, N52_HC_, and S54_HC_ form a hydrogen bond network with E484_RBD_. The side chains of Y91_LC_ and S95_LC_ hydrogen bond with backbones of G485_RBD_ and F486_RBD_. In addition, Y30_LC_, Y32_LC_, Y91_LC_, and Y100j_HC_ also form a hydrophobic pocket to hold F486_RBD_ (Figure S3B), with π-π stacking observed between Y30_LC_ and F486_RBD_ (Figure 3A right).

**Figure 3.**
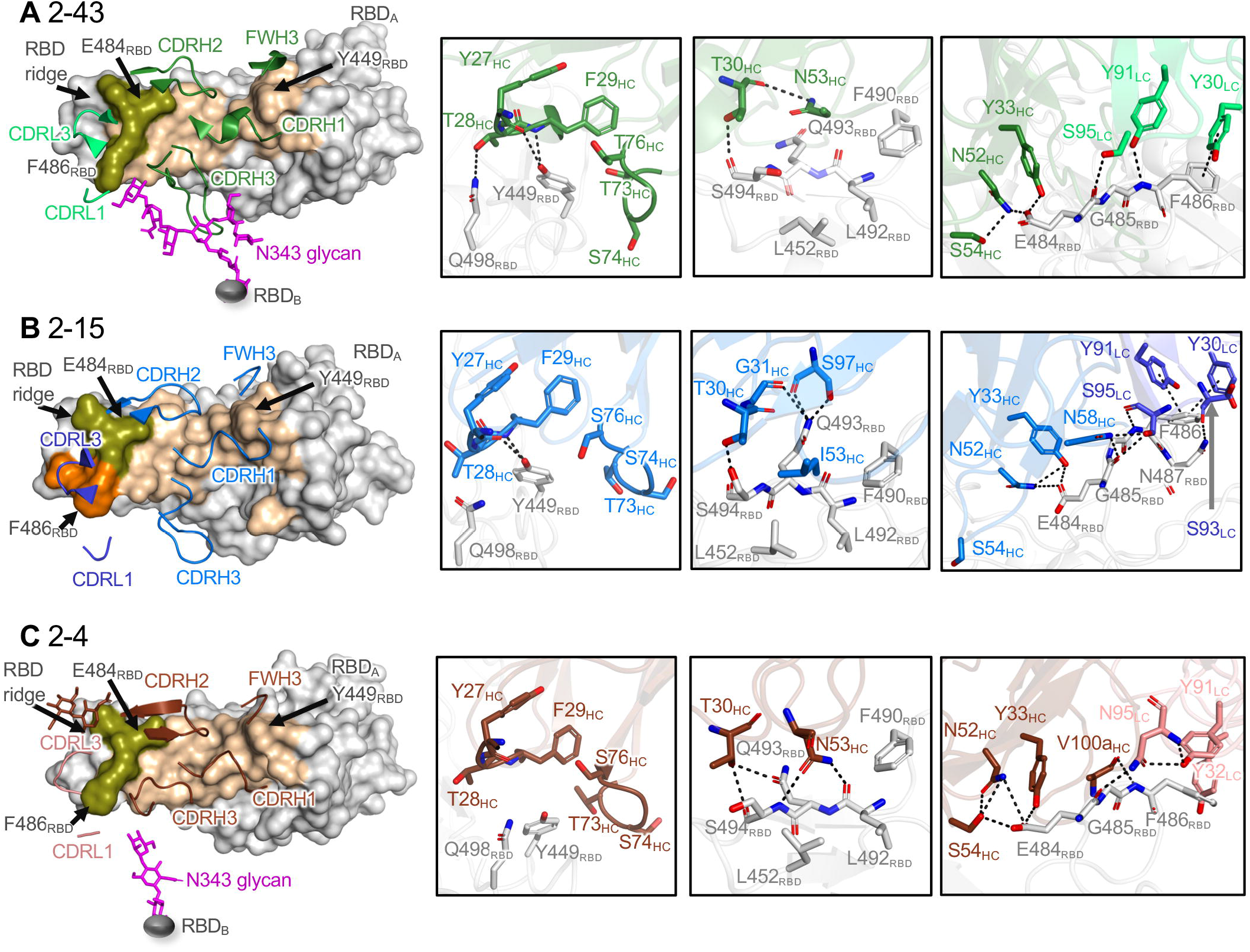
Common epitope-paratope interactions define a VH1-2 antibody class. (A) Overview of the 2-43 epitope (left panel) and close-up view of the hydrogen bond networks between 2-43 and the SARS-CoV-2 RBD (right three panels). The RBD epitope recognized by 2-43 heavy (forest) and light (limegreen) chains are colored wheat and orange respectively(RBD_A_, light gray). Epitope residues interacting with both heavy and light chains are colored lemon. 2-43 also binds to N343 glycan (magenta) from a neighboring RBD (RBD_B_, dark gray). Hydrogen bonds and π-π stacking are shown as black dashed lines. (B) Overview of the epitope of 2-15 (left panel) and close-up view of the hydrogen bond networks and π-π stackings between 2-15 and the SARS-CoV-2 RBD (right three panels). 2-15 heavy and light chains are colored marine and blue respectively. Epitope residues interacting with both heavy and light chains are colored lemon. (C)Overview of the epitope of 2-4 (left panel) and close-up view of the hydrogen bond networks between 2-4 and the SARS-CoV-2 RBD (right three panels). The heavy and light chains of 2-4 are colored chocolate and salmon respectively. Epitope residues interacting with both heavy and light chains are colored lemon. See also Figure S3.

The crystal structure of 2-15 complexed with SARS-CoV-2 RBD revealed 2-15 to use similar heavy and light chain regions to recognize the RBM (Figure 3B left panel). Similar to 2-43, 2-15 forms convergent hydrogen bonds to Y449_RBD_, S494_RBD_, E484_RBD_, and G485_RBD_ through residues Y27_HC_, F29_HC_, T30_HC_, Y33_HC_, N52_HC_, and S95_LC_ (Figure 3B right three panels). 2-15 Y30_LC_ and Y91_LC_ engage in π-π stackings with F486_RBD_. Nonetheless, 2-15 interactions different from 2-43 are also observed at the RBM flat and ridge regions. The side chain of 2-15 I53_HC_ interacts with a hydrophobic pocket formed by residues L452_RBD_, F490_RBD_, L492_RBD_, Q493_RBD_, and S494_RBD_– interactions that are not observed in 2-43 which instead includes germline residue N53_HC_ (Figure 3B third panel). Additional hydrogen bonds are formed between G31_HC_ and S97_HC_ and Q493_RBD_ and between N58_HC_ and E484_RBD_ (Figure 3B right two panels).

Compared to 2-43 and 2-15, the published VH1-2 antibodies 2-4, S2M11, and C121 showed similar RBM approach angle and binding mode (Figure 3C, 3D, S3C, and S3D). Typically, interactions between the VH1-2 antibodies and the RBD buried a total of about 1300-1600Å^2^ BSA (RBD_A_ column in Figure S3C), with antibody heavy chains contributing about 80% of the paratope. For all antibodies, shared paratope residues are observed in FWH1, CDRH1, CDRH2, FWH3, CDRL1, and CDRL3 (Figure S1A and S1C). FWH1, CDRH1, and CDRH2 residues form conserved hydrogen bond networks with Y449_RBD_, E484_RBD_, and Q493. Thus, these VH1-2 encoded residues form a module for RBM recognition. Despite these antibodies having different light chain gene origins, convergent tyrosine residues (Y30_LC_, Y32_LC_, and Y91_LC_) in CDRL1 and CDRL3 pack against F486_RBD_ (Figure S3B), which constitutes another module for RBD recognition. Because each antibody has a unique CDRH3, no conserved polar or hydrophobic interaction is observed (Figure 3 and S3B-S3D), albeit CDRH3 contributes the most BSA among the CDRs except in 2-15 (Figure S3A). Altogether, the germline gene residues from the VH1-2 gene as well as light chain V genes anchor the antibodies to the RBM in a similar mode with heavy chain V region playing a dominant role in determining the mode of recognition.

### The VH1-2 antibody class and similarity to other SARS-CoV-2 RBD targeting antibodies

To gain an overall understanding of similarity in binding orientations of the VH1-2 antibodies, we superposed RBD and antibody complexes and calculated pairwise root mean square deviation (RMSD) to compare relative binding orientations of the six VH1-2 antibodies (2-43, 2-15, H4, 2-4, C121, and S2M11) as well as 31 additional RBD-targeting antibodies originated from other VH genes. Overall, the VH1-2 antibodies have similar binding orientations (Figure 4A). Clustering of the 37 antibodies using pairwise RMSDs showed that the six VH1-2 antibodies group together (Figure S4A). The six VH1-2 antibodies also have highly overlapping heavy and light chain approach angles and epitopes (Figure 4B and S4B), which overlap with ACE2 binding site (Figure S4D). The similarity in genetic origin, details of their interactions with RBD (unavailable for H4), and angle of approach, suggests that these six antibodies form a VH1-2 antibody class.

**Figure 4.**
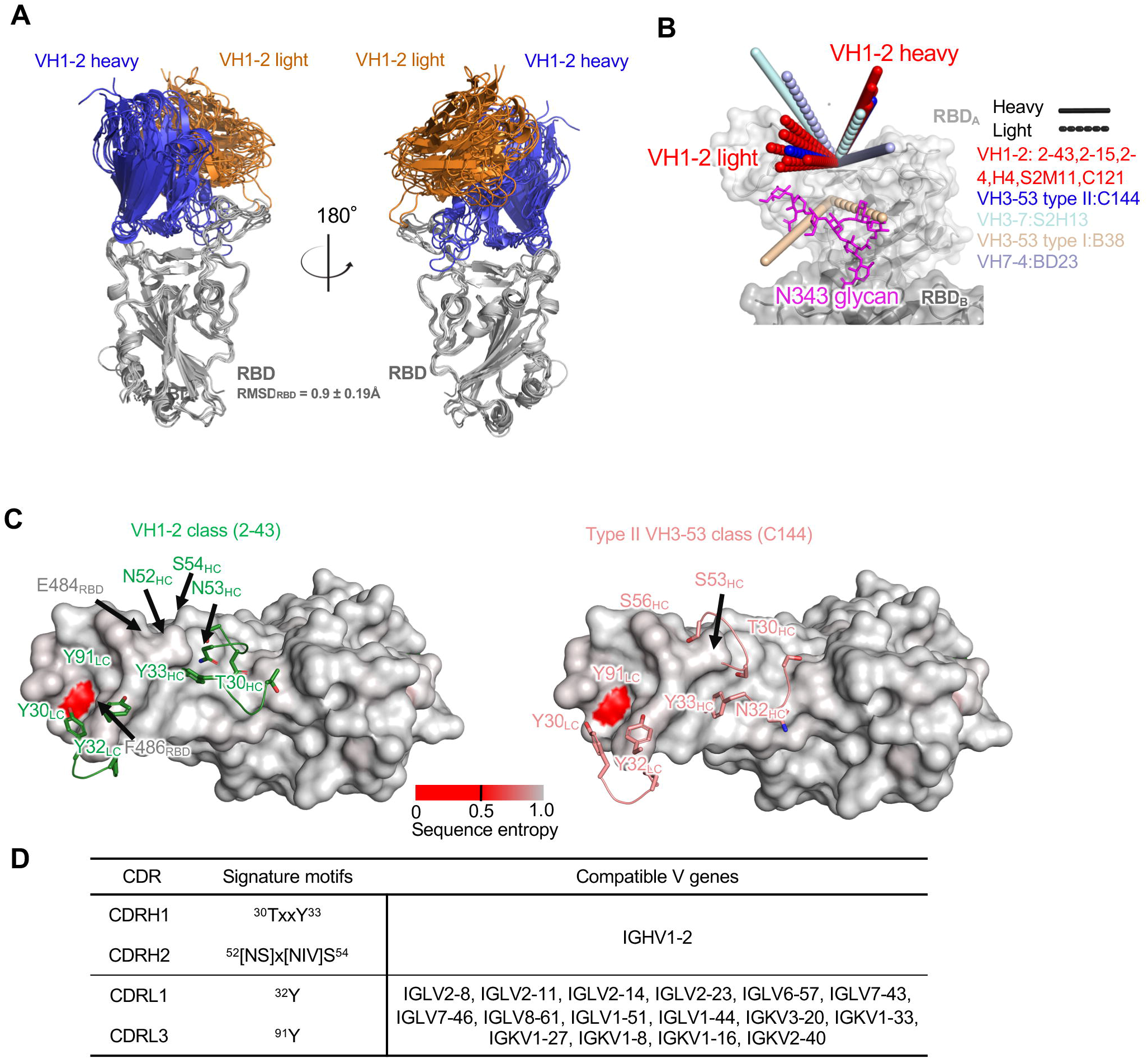
The VH1-2 antibody class. (A) Overview of modes of RBD recognition by the VH1-2 class antibodies 2-43, 2-15, 2-4, H4, S2M11, and C121. The RBD and antibody complexes are superimposed on RBDs. (B) Approach angles of SARS-CoV-2 RBD-targeting antibodies originated from different VH genes. Members of the VH1-2 antibody class have similar approach angles. The VH3-53 class antibody C144 has an angle of approach similar to the VH1-2 antibody class. (C) Overview of VH1-2 signature motifs (green) and type II VH3-53 class (orange) on RBD surface. The RBD is colored with conservation score calculated from circulating SARS-CoV-2 strains. 2-43 structure is used to generate the conservation plot. (D) Signature motifs and compatible germline V genes of the VH1-2 antibody class. See also Figures S1 and S4.

The structural, recombination, and somatic hypermutation analyses (described below in the next section) presented here revealed critical residues that determine the specificity and binding affinity of the VH1-2 class antibodies. We define residues forming conserved side chain polar and/or hydrophobic interactions as genetic signatures. The heavy chain signatures include a T30-x-x-Y33 motif in CDRH1 and a [NS]52-x-[NIV]-S54 motif (x represents any amino acid) in CDRH2 (Figure 4C), with no conserved motif observed in CDRH3 (Figure S1A). Searches of germline gene databases using this signature showed only alleles of the VH1-2 gene to match both motifs (Figure 4E), suggesting that the antibody class is likely to be strictly VH1-2-restricted.

For light chain, the side chains of Y32_LC_ and Y91_LC_ interact with F486_RBD_ (Figure 3 and S3B). In 2-43, 2-4, and S2M11, Y32_LC_ also interacts with N343 glycan from adjacent RBDs (described below in the quaternary recognition section). Because the light chain interaction-signatures can be found in many germline genes (Figure 4D and S4C), the VH1-2 antibody class may not be restricted with respect to light chain origin gene.

To understand the conservation of the VH1-2 antibody epitope, we calculated the conservation score for each RBD residue and observed that the flat region is highly conserved in the natural SARS-CoV-2 reservoir. The four RBD residues critical for VH1-2 antibody recognition, Y449_RBD_, E484_RBD_, F486_RBD_, and Q493_RBD_, showed low mutation frequencies (approximately 0, 3, 7, and 7 per 10,000 sequences, respectively). Nonetheless, SARS-CoV-2 strains with mutations at the four positions are resistant to C121 and S2M11 of the VH1-2 antibody class (Tortorici et al., 2020, Barnes et al., 2020a).

In addition, the VH1-2 antibodies show high similarity in approach angle and epitope to antibody C144 (Figure 4C and S4A), a type II VH3-53/-66 antibody. However, C144 uses different CDRH1 and CDRH2 motifs for interacting with the conserved residues in the flat region and ridge of RBD (e.g. the N32-Y33 and [T/S]54-G-G-[T/S]57 motifs in the VH3-53/-66 antibodies) (Figure 4C) (Wu et al., 2020a, Barnes et al., 2020a). The CDRH2 of C144 shifts towards the RBD ridge, perhaps because the CDRH2s of the VH3-53/-66 genes are one residue shorter than the VH1-2 gene. Nonetheless, both antibody classes use similar light chain genes including VL2-14 and VL2-23, which provide key Tyr residues that interact with F486_RBD_.

### Somatic hypermutations and avidity improve antibody potency

To understand the effects of somatic hypermutation (SHM) on binding affinity and neutralization potency, we reverted SHMs in the paratope regions of 2-43, 2-15, and 2-4 to their germline residues individually and in combination. 2-43 only has one SHM in the paratope region, S76T_HC_ (Figure S1A), the reversion of which reduces the IgG apparent binding affinity and pseudovirus neutralization potency by about 2-fold and 5-fold respectively (Figure 5A, 5B, and S4E). For 2-15, three paratope SHMs were observed: N53I_HC_, G55D_HC_, and Y32F_LC_. The reversion of N53I_HC_ and Y32F_LC_ individually reduced binding affinity by 16-fold and 5-fold respectively (Figure 5A) as well as neutralization potency by 217-fold and 42-fold respectively (Figure 5B). The results suggested that the N53I_HC_ and F32Y_HC_ mutations are critical for the affinity maturation of 2-15. Structural analysis showed that the N53I_HC_ mutation allows the Ile side chain to fit a hydrophobic pocket on RBD (Figure 3B and S5A). A convergent mutation is also observed in S2M11 (N53I_HC_) and C121 (N53V_HC_). We then introduced the N53I_HC_ mutation to 2-43 and 2-4 and observed significant improvements of both binding affinity and neutralization potency (Figure 5). These results suggested that mutation to a hydrophobic residue at 53_HC_ can be used to improve members of the VH1-2 antibody class. The crystal structure further showed that 2-15 32_LC_ does not interact with RBD, but instead locates between W100g_HC_ and Y91_LC_, which interact with RBD (Figure S3B). This implies that the Y32F_LC_ SHM may alter the interaction with RBD indirectly. For 2-4, the SHM A60T_HC_ generates an N-glycosylation site at position N58_HC_. The N58_HC_ glycan interacts with the RBD ridge (Figure 3C). The reversion of this SHM reduces binding affinity and neutralization by about 5-fold and 9-fold respectively (Figure 5). In summary, somatic hypermutation analysis revealed that the precursors of these antibodies could bind antigens with nanomolar apparent binding affinity, suggesting that the precursors of these antibodies can be activated efficiently by the SARS-CoV-2 spike protein. Nonetheless, SHMs significantly improve both the binding affinity and the neutralization potency of these antibodies. Because the observed SHMs are frequently generated by the somatic hypermutation machinery (Figure S1A), we anticipate that requirements for somatic hypermutations are unlikely to present a significant barrier for affinity maturation of this antibody class.

**Figure 5.**
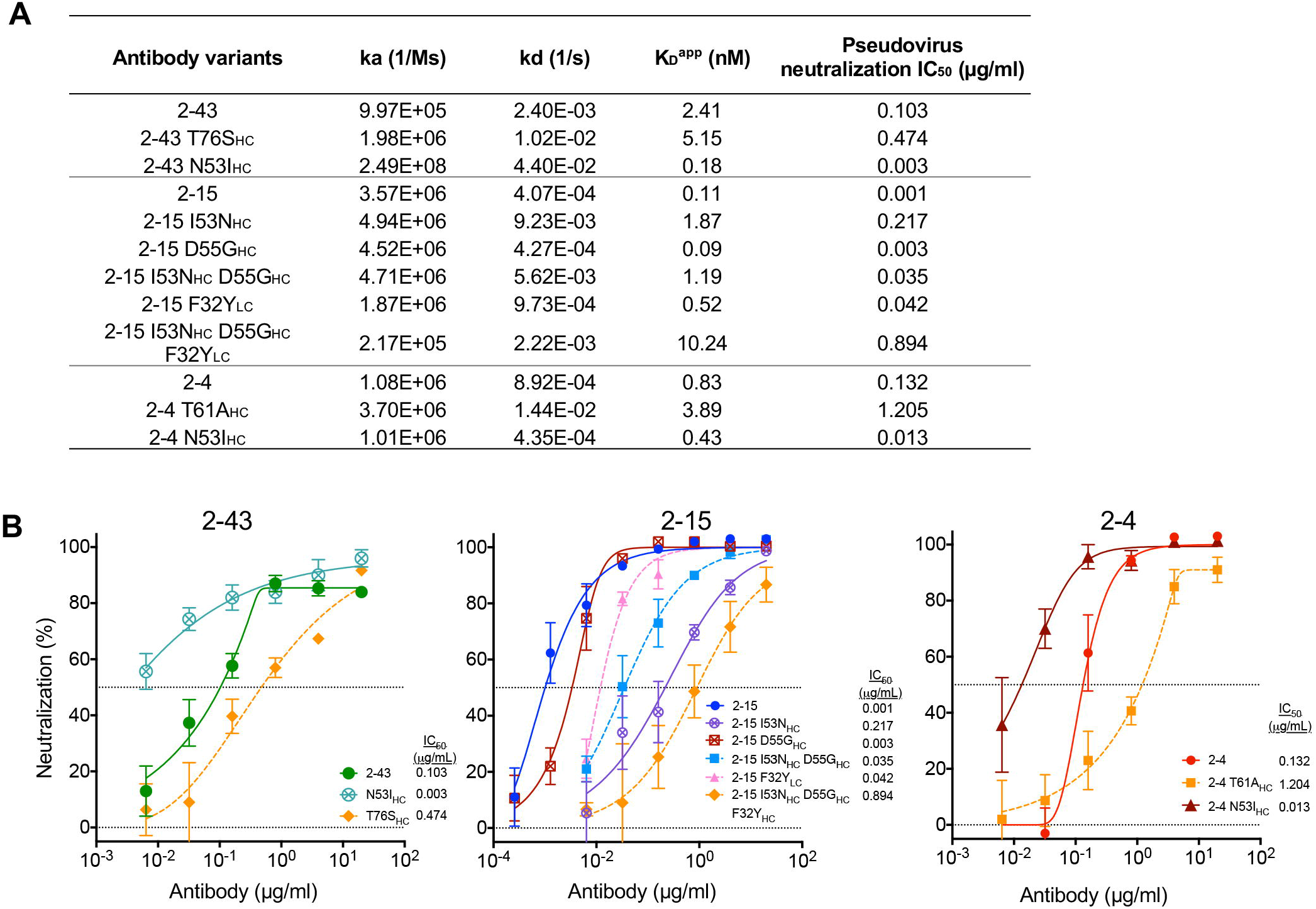
N53I mutation improves many VH1-2 class members. (A) Apparent binding affinities of 2-43, 2-15, and 2-4 lgGs and revertants. Somatic hypermutations improve binding affinity of the three antibodies. N53I _Hc_ was introduced to 2-43 and 2-4 and showed significant increased binding affinity. (B) Pseudovirus neutralization profiles of 2-43, 2-15, 2-4 and their somatic hypermutation revertants. See also Figures S1, S3, and S5.

In addition, we evaluated effect of avidity on antibodies 2-43, 2-15, and 2-4. Comparison of neutralization potency between IgGs and Fabs revealed that the Fabs of 2-43, 2-15, and 2-4 have potencies of about 140-, 95-, and 14-fold lower than their IgG forms respectively (Figure S5C), suggesting that avidity effects are critical for achieving high neutralization potency.

### Recognition of quaternary epitopes modulates spike conformation

Our previous study with a low resolution cryo-EM structure showed that 2-43 recognizes a quaternary epitope (Liu et al., 2020). However, details on the quaternary interactions and their functional relevance have not yet been characterized. Here, high resolution cryo-EM reconstructions revealed atomic-level interactions between 2-43 and the quaternary epitope. Overall, 2-43 interacts comprehensively with N343 glycan as well as helix 364-371 from an adjacent ‘down’ RBD protomer (RBD_B_) (Figure 6A and S6A), burying a total of 999Å BSA (Figure S3A). The quaternary interaction is predominantly mediated by the long CDRH3, which is held by the two branches of the N343 glycan, the structure of which is highly flexible and has not been observed in previous studies. The 2-43 CDRH3 forms two hydrogen bond networks to stabilize N343 glycan (Figure 6A). The tip of one N343 glycan branch is further stabilized by hydrogen bonding with 2-43 Y32_LC_ and the ridge of RBD_A_ (N487_RBD_ and Y489_RBD_) (Figure S6A). In addition, the tip of the elongated CDRH3 interacts with helix 364-371 of RBD_B_ (Figure 6A). Altogether, 2-43 forms strong quaternary interactions with the adjacent ‘down’ RBD, which is critical for locking RBDs in the all ‘down’ conformation. In contrast to 2-43, the cryo-EM density maps of the 2-15 and H4 spike complexes revealed no quaternary contacts (Figure 2D, S2L, and 6B).

**Figure 6.**
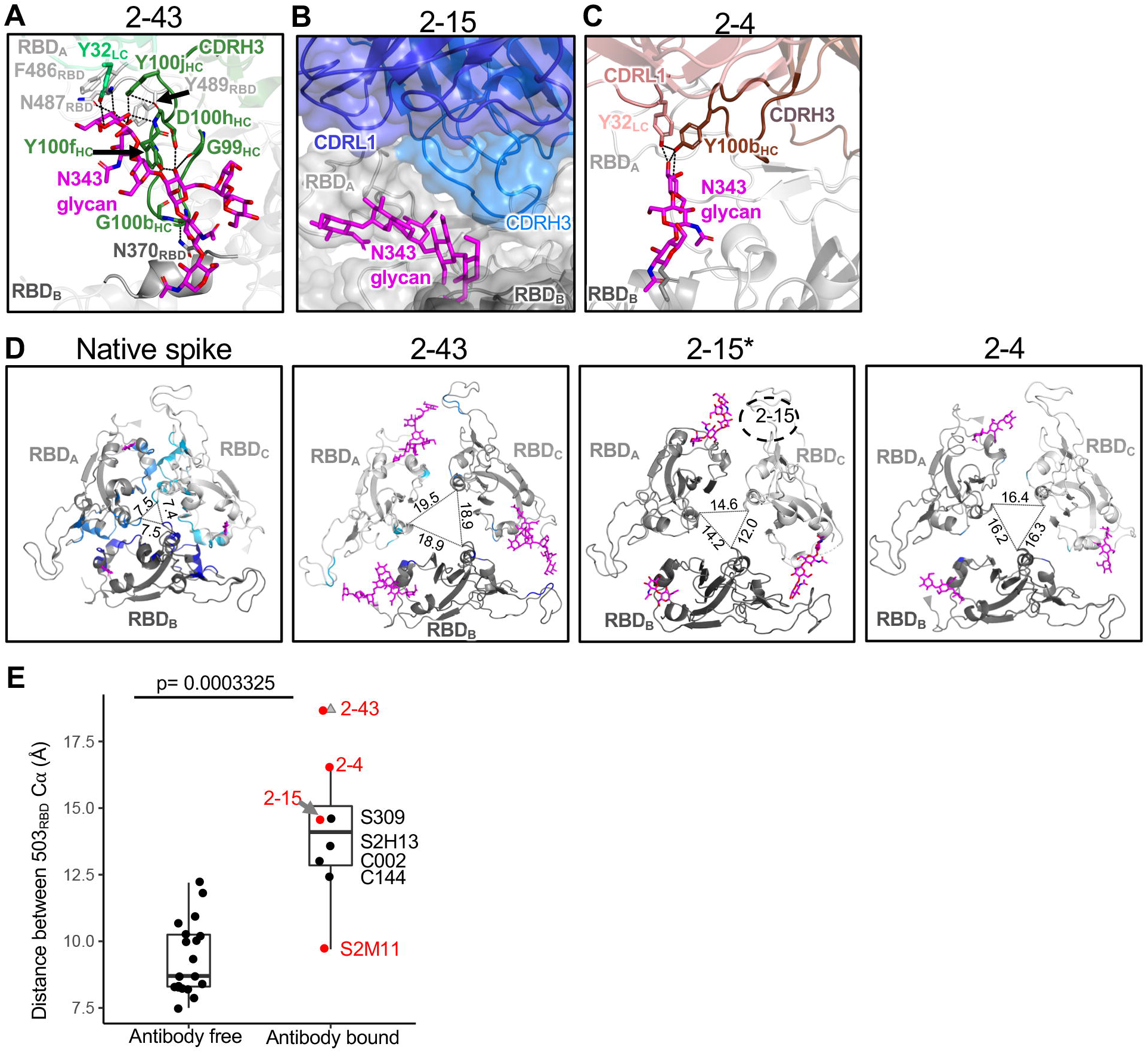
Subset of the VH1-2 antibody class uses an elongated CDRH3 to recognize glycan N343 from a neighboring RBD, a quaternary interaction that expands the SARS-CoV-2 spike. (A) Close-up view of the quaternary epitope of 2-43. The 2-43 CDRH3 forms two hydrogen bond networks (black dashed lines) with an adjacent RBD (RBD_B_) N343 glycan and also interacts with helix 364-371 of RBD _B_. (B) Close-up view of potential quaternary interactions between 2-15 and the neighboring RBD (RBD_B_) . 2-15 reconstructed from cryo-EM data shows minor quaternary interaction. (C) Close-up view of the interaction between 2-4 and N343 glycan of the neighboring RBD. (D) Distance between antibody free and antibody bound trimeric RBDs. Antibody binding induces significantly larger expansion of the trimeric RBDs. The three RBDs were colored gray. Residues at RBD interfaces were colored cyan, light blue, and blue for protomers RBD_A_, RBD_B_, and RBDc respectively. The distances between RBD protomers were measured using Cα of position 503. Note: due to low resolution, the interface residues between RBDs were not shown for 2-15 bound spike. 2-15 (dashed circle) binds to protomer RBDc. (E) Distance of Cα of position 503 between the trimeric ‘down’ RBDs. Antibody-free spikes have a shorter distance than the antibody-bound spikes. For the boxplots, the middle lines are medians. The lower and upper hinges correspond to the first and third quartiles respectively. Mann Whitney U Test was used. Outliers are shown as gray triangles. See also Figures S2, S3, and S6.

Comparison of quaternary interactions among the five VH1-2-derived antibodies revealed two structural groups. Group 1 includes 2-43, 2-4, and S2M11, which bind predominantly ‘down’ RBDs. Group 2 includes 2-15, H4, and C121, which bind both ‘up’ and ‘down’ or only ‘up’ RBDs. For group 1, the quaternary recognition is mediated mostly by CDRH3, followed by CDRL1 and CDRL2 (Figure 6C and S3A). Differences between each antibody-specific CDRH3 determines that each antibody has a unique quaternary epitope (Figure 6A, 6B, 6C, S3B, and S6B). Different from S2M11, the quaternary interaction is indispensable for 2-43 which binds to spike trimer but not isolated RBD (Liu et al., 2020). The quaternary epitope of 2-4, only part of which appears to be visible in the cryo-EM map (Liu et al., 2020) comprises only N343 glycan from the adjacent ‘down’ RBD protomer (Figure 6C), likely because 2-4 has a short CDRH3. A hydrogen bond network is observed between CDRH3 Y100b_HC_, CDRL1 Y32_LC_, and N343 glycan of the adjacent RBD. The quaternary epitope of S2M11 includes N343 glycan and helices 339-343 and 364-371 of the adjacent RBD (Figure S6B). Altogether, accommodation of N343 glycan from the adjacent ‘down’ RBD is a common feature of quaternary recognition by group 1 antibodies. The diverse CDRH3s of the group 1 antibodies form a module for quaternary recognition. In contrast, group 2 antibodies 2-15 and C121 show minor interactions with neighboring ‘down’ RBDs (Figure 2C, 6B, S2R, and S3A), which may explain why they do not lock the trimeric RBDs in an all ‘down’ conformation.

To understand whether antibody binding induces conformation change in the SARS-CoV-2 spike protein, we compared the antibody-bound and ligand-free spike structures in the closed (all ‘down’) prefusion conformation. These comparisons showed that the binding of the VH1-2 antibodies (except S2M11), as well as other RBD-directed antibodies leads to significantly larger distances and less contacts between the trimeric ‘down’ RBDs (Figure 6D and 6E). Each of these antibodies plays a role in bridging the interactions between RBDs as well as reorienting the conformation of N343 glycan. However, antibodies like 2-15, which does not form strong interactions with the adjacent RBD, may disassemble the spike when 3 Fabs bind to the RBDs (Figure S2J and S2K). In addition, superposition of 2-15 on the ‘down’ RBD in ligand-free spike revealed the light chain to significantly clash with an adjacent ‘up’ RBD (Figure S6D), suggesting members of the VH1-2 antibody class to either bind only ‘down’ RBDs when a neighboring ‘down’ RBD is available or push the neighboring ‘up’ RBD away to bind ‘down’ RBDs.

Consistent with the first mechanism, we did not observe a neighboring ‘up’ RBD adjacent to 2-15 and 2-4 bound ‘down’ RBDs in the cryo-EM data. In contrast, C121 adopts the second mechanism through light chain interaction with the adjacent ‘up’ RBD (Figure S6D) (Barnes et al., 2020a). In summary, the quaternary interaction and angle of approach determine that the VH1-2 antibody class modulates SARS-CoV-2 spike conformation when binding to a ‘down’ RBD.

## Discussion

In this study, we determined structures of three nAbs: 2-43, 2-15, and H4, which revealed a VH1-2 antibody class with a common RBD-binding mode and similar angle of approach. The VH1-2 antibody class uses two modules for spike recognition with the VH1-2 gene encoded module for recognition of RBD and CDRH3 module for quaternary recognition. The VH1-2 antibody class has little or no restraint on CDRH3 length. The prevalence of the VH1-2 antibody class in numerous donors (Liu et al., 2020, Wu et al., 2020b, Hansen et al., 2020, Robbiani et al., 2020, Rogers et al., 2020, Brouwer et al., 2020, Ju et al., 2020, Zost et al., 2020, Kreer et al., 2020, Tortorici et al., 2020) suggests it to be a common component of the effective antibody response to SARS-CoV-2 that can include highly potent neutralizing antibodies.

For some members of the VH1-2 class, recognition of a quaternary epitope can lock RBDs in the ‘down’ conformation, providing an additional mechanism to achieve high neutralization potency. The quaternary epitopes of the VH1-2 class include the N343 glycan and helix 364-371 of an adjacent RBD. A similar quaternary epitope is recognized by the type II VH3-53/-66 class antibody C144 (Barnes et al., 2020a). Thus, this quaternary epitope appears to represent a supersite at which antibodies lock the RBDs in the all ‘down’ conformation. We also observe that the quaternary interaction can induce distinct conformational changes of the trimeric ‘down’ RBDs. The quaternary interaction observed in S2M11 does not significantly alter the distance between spike protomers (Figure 4E) (Tortorici et al., 2020). In contrast, the quaternary interactions of 2-43 and 2-4, mediated predominantly by CDRH3, increase the distance of the trimeric RBDs (Figure 4). For 2-15, the cryo-EM structure of one Fab bound spike revealed a CDRH3 conformation moving away from the adjacent ‘down’ RBD to avoid steric clash. Despite this, 2-15 also increases the distance between RBDs, which may be the result of mobile quaternary contacts that cannot be observed in the low resolution cryo-EM reconstruction. We anticipate that the interactions between RBDs could be further weakened when all three protomers are bound by 2-15, which may be the cause of the observed spike disassembly by 2-15 (Figure S2J and S2K). The disassembly of spike may be another mechanism for 2-15 to achieve ultrapotent neutralization. In addition, the crystal structure of 2-15 in complex with isolated RBD showed 2-15 to have another stable binding mode in the absence of a quaternary contact. We hypothesize that this binding mode may be observed when 2-15 binds to ‘up’ RBDs and cannot form quaternary interactions.

The structural information we present also provides clues for further optimization of the VH1-2 antibody class. Despite members of the class achieving high potency with germline gene mediated interactions, we observe that somatic hypermutations can further improve this class significantly. The conserved mode of RBD recognition implies that members of the antibody class will be sensitive to similar viral escape mutations (e.g. C121 is sensitive to E484K_RBD_ and Q493R_RBD_) (Barnes et al., 2020a, Tortorici et al., 2020).

## Acknowledgements

Cryo-EM data collection was performed at the National Center for Cryo-EM Access and Training and the Simons Electron Microscopy Center located at the New York Structural Biology Center, supported by the NIH Common Fund Transformative High Resolution Cryo-Electron Microscopy program (U24 GM129539) and by grants from the Simons Foundation (SF349247) and NY State Assembly. Some of this work was also performed at the Columbia University Cryo-Electron Microscopy Center. Data analysis was performed at the National Resource for Automated Molecular Microscopy (NRAMM), supported by the NIH National Institute of General Medical Sciences (GM103310). We thank D. Neau, S. Banerjee, and Igor Kourinov for help with synchrotron data collection conducted at the APS NE-CAT 24-ID-C beamline, which is supported by National Institutes of Health (NIH) P41 GM103403; use of NE-CAT at the Advanced Photon Source was supported by the US Department of Energy, Basic Energy Sciences, Office of Science, under contract number W-31-109-Eng-38. Support for this work was provided by the Intramural Research Program of the Vaccine Research Center, National Institute of Allergy and Infectious Diseases (NIAID), National Institutes of Health (NIH). This study was supported by Samuel Yin, Pony Ma, Peggy & Andrew Cherng, Brii Bioscieces, Jack Ma Foundation, JBP Foundation, Carol Ludwig, and Roger & David Wu.

## Author Contributions

M.R., and G.C. determined cryo-EM structures of 2-43, 2-15, and H4; Y.G. performed bioinformatics analyses; E.R. solved crystal structure of 2-15; L.L. performed SPR; P.W. performed neutralization assessment; J.Y. produced antibodies; D.D.H. supervised antibody production, SPR, and neutralization assays; P.D.K. supervised reagents, L.S. supervised cryo-EM and crystal structure studies; Z.S supervised informatics studies. L.S. and Z.S. oversaw the project and –with M.R., Y.G., E.R., L.L., P.W., J.Y., and B.Z.– wrote the manuscript, with all authors providing revisions and comments.

## STAR⍰METHODS

Detailed methods are provided in the online version of the paper and include the following:

- KEY RESOURCES TABLE
- RESOURCE AVAILABLITY
  - Lead contact
  - Materials availability
  - Data and code availability

- EXPERIMENTAL MODEL AND SUBJECT DETAILS
  - Cell lines

- METHOD DETAILS
  - Expression and Purification of SARS-CoV-2 Spike and RBD Proteins
  - Antibody production and mutagenesis
  - Production of Fab from IgG
  - Binding Affinity Measurements by Surface Plasmon Resonance
  - Pseudovirus neutralization assays
  - Antibody gene assignments and genetic analyses
  - Delineation of sequence signatures
  - Analysis of antibody repertoire data (NGS)
  - Frequencies of sequence signatures of the VH1-2 and VH3-53/-66 antibody classes
  - Cryo-EM data collection and processing
  - Cryo-EM model building
  - X-ray data collection, structure solution, model building and refinement

- QUANTIFICATION AND STATISTICAL ANALYSIS

## KEY RESOURCES TABLE

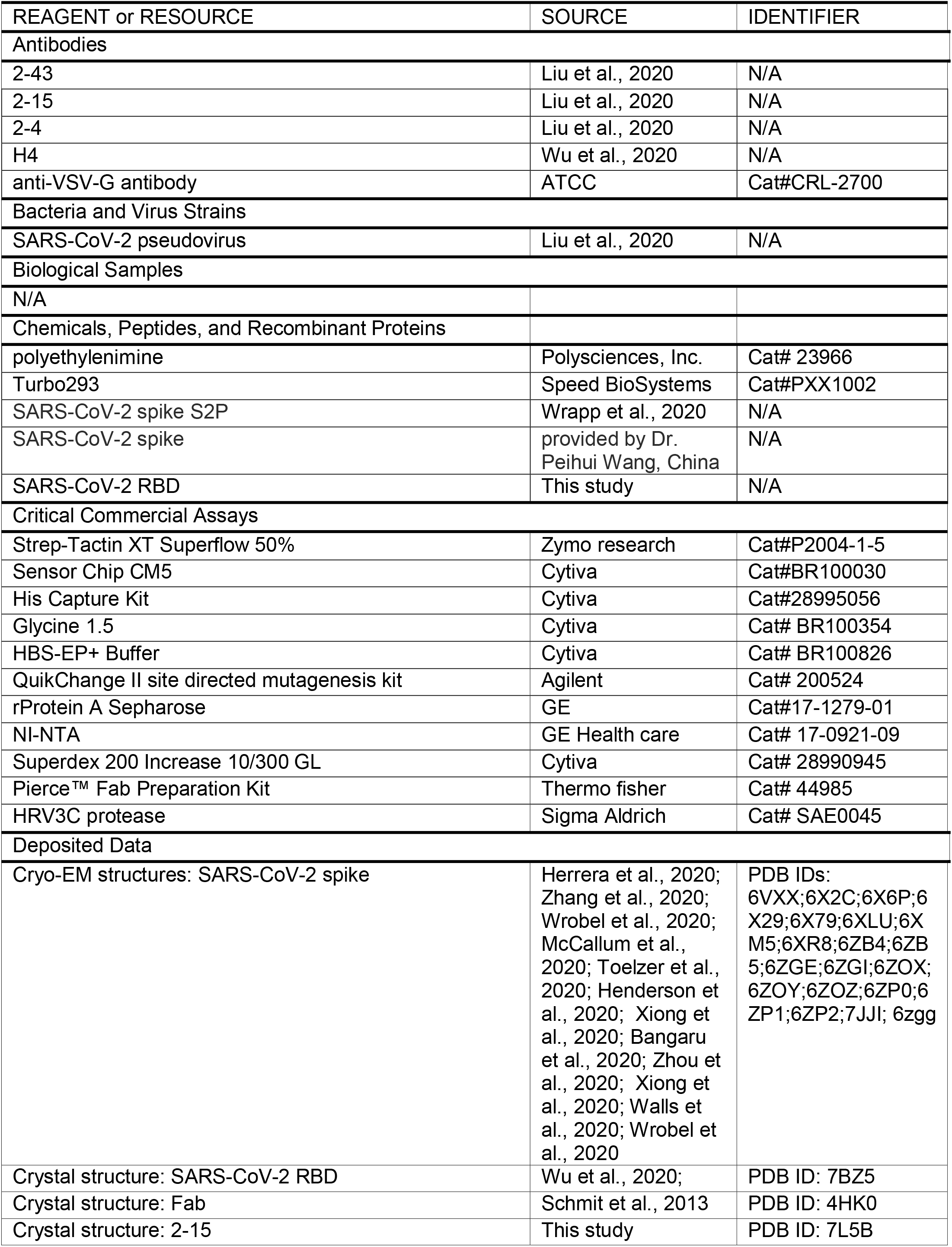
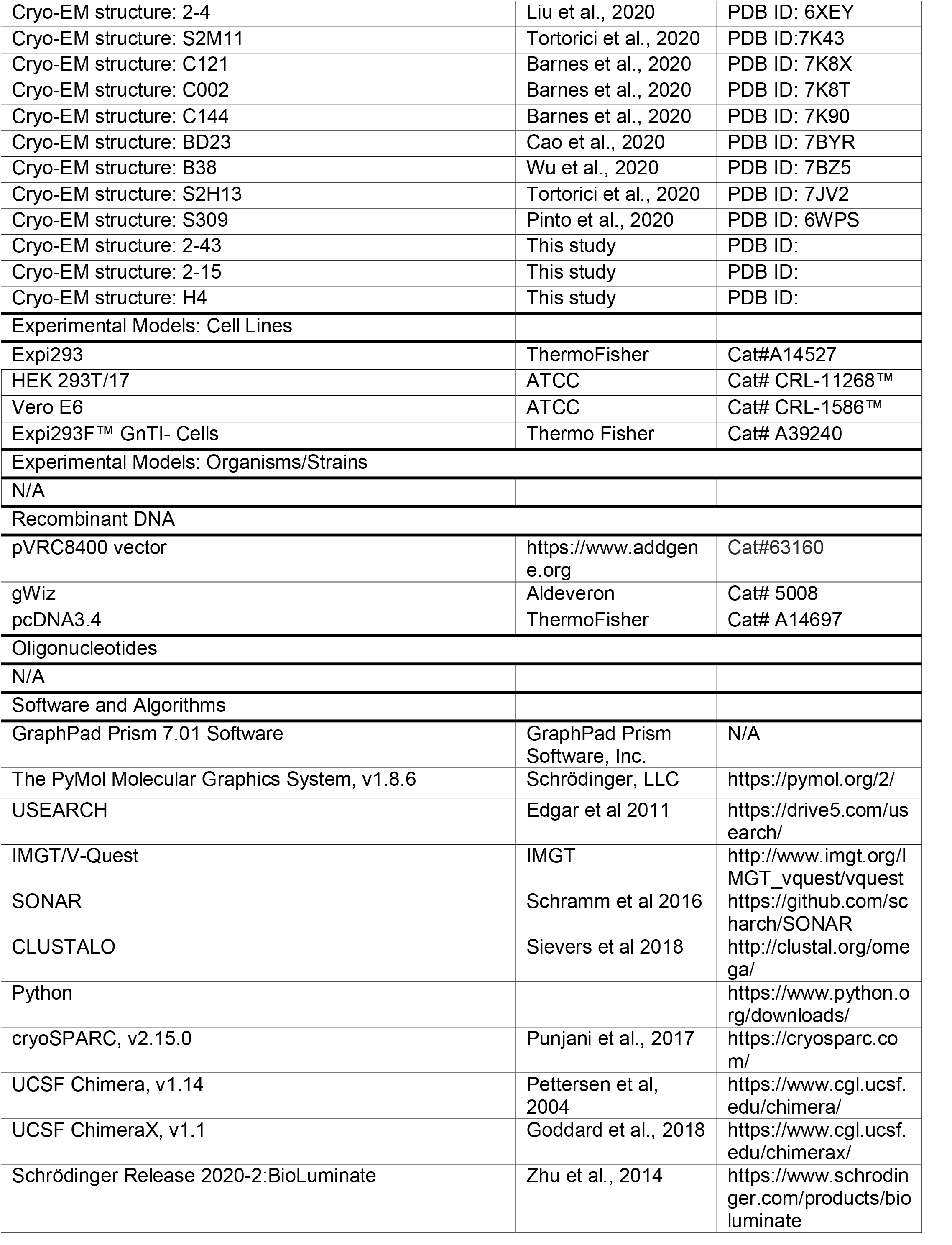
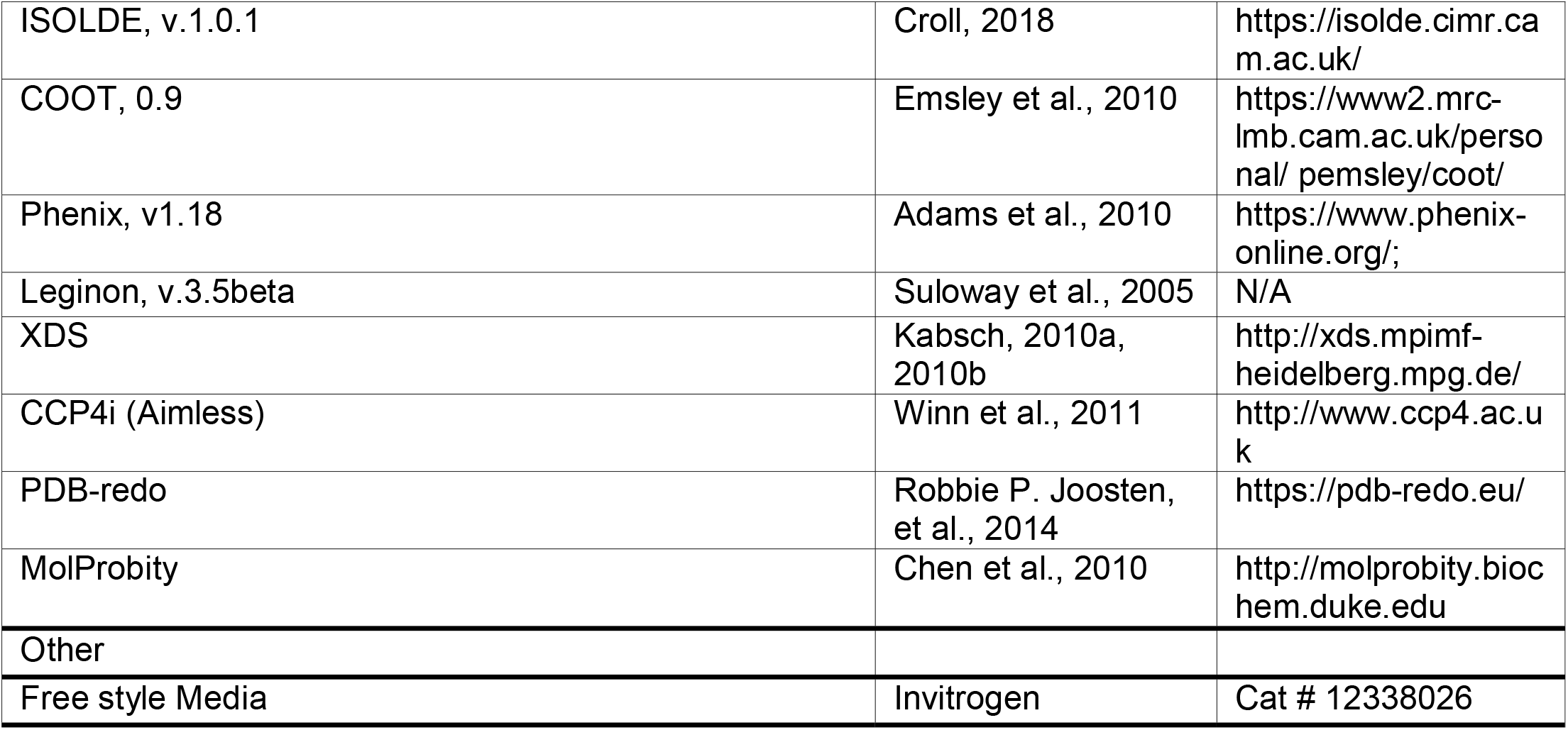

## RESOURCE AVAILABILITY

### Lead Contact

Further information and requests for resources and reagents should be directed to and will be fulfilled by the Lead Contact, Zizhang Sheng.

### Materials Availability

Expression plasmids generated in this study for expressing SARS-CoV-2 proteins and antibody mutants will be shared upon request.

### Data and Code Availability

Cryo-EM maps and fitted coordinates are in the process of being deposited to the EMDB and RCSB respectively. The accession numbers of the published next-generation sequencing data can be found in the Key Resources Table.

## EXPERIMENTAL MODEL AND SUBJECT DETAILS

### Cell lines

Expi293 cells and Expi293F™ GnTI-Cells were from ThermoFisher Scientific Inc (ThermoFisher, cat#A14527 and cat# A39240 respectively). HEK 293T/17(cat# CRL-11268(tm)) and Vero E6 cells (cat# CRL-1586™) were from ATCC.

## METHOD DETAILS

### Expression and Purification of SARS-CoV-2 Spike and RBD Proteins

The mammalian expression vector that encodes the ectodomain of the SARS-CoV-2 S trimer was kindly provided by Jason McLellan (Wrapp et al., 2020). SARS-CoV-2 S trimer expression vector was transiently transfected into Expi293™ cells using 1 mg/mL of polyethylenimine (Polysciences). Five days post transfection, the S trimer was purified using Strep-Tactin XT Resin (Zymo Research).

The SARS-CoV-2 RBD (residues 331-528), was cloned into the pVRC-8400 mammalian expression plasmid, with a C-terminal 6X-His-tag and an intervening HRV-3C protease cleavage site. Expression vector was transiently transfected into Human Embryonic Kidney (HEK) 293 GnTI-Freestyle cells suspension culture in serum-free media (Invitrogen) using polyethyleneimine (Polysciences). Media was harvested 4 days after transfection and the secreted protein purified using Ni-NTA IMAC Sepharose 6 Fast Flow resin (GE Healthcare) followed by size exclusion chromatography (SEC) on Superdex 200 (GE Healthcare) in 10 mM, Tris pH 8.0, 150 mM NaCl. Peak fractions containing RBD were pooled and HRV-3C protease (Thermo fisher) was added in a mass ratio 1:100 relative to RBD, followed by incubation for 24 h at 20°C, to remove the C-terminal histidine-tag. The protein solution was again passed through the NI-NTA column to remove the His-tag and any traces of uncleaved protein. Protein purity was analyzed by SDS-PAGE and buffer exchanged to SEC buffer and concentrated to ∼5 mg/mL and used for crystallization experiments.

### Antibody production and mutagenesis

For each antibody, the variable genes were optimized for human cell expression and synthesized by GenScript. VH and VL were inserted separately into plasmids (gWiz or pcDNA3.4) that encoding the constant region for heavy chain and light chain respectively. Monoclonal antibodies were expressed in Expi293 (ThermoFisher, A14527) by co-transfecting heavy chain and light chain expressing plasmids using polyethylenimine (PEI, Linear, MV∼25,000, Polysciences, Inc. Cat. No. 23966) and culture in 37.0C degree shaker at 125RPM and 8% CO2. Supernatants were collected on day 5, antibodies were purified by rProtein A Sepharose (GE, 17-1279-01) affinity chromatography.

For H4, the antibody expression constructs were synthesized (Gene Universal Inc, Newark DE) and subcloned into corresponding pVRC8400 vectors. To express the antibodies, equal among of heavy and light chain plasmid DNA were transfected into Expi293F cells (Life Technology) by using Turbo293 transfection reagent (Speed BioSystems). The transfected cells were cultured in shaker incubator at 120 rpm, 37 °C, 9% CO2 for 5 days. Culture supernatants were harvested and purified over Protein A (GE Health Science) resin in columns. Each antibody was eluted with IgG elution buffer (Pierce), immediately neutralized with one tenth volume of 1M Tris-HCL pH 8.0. The antibodies were then buffer exchanged in PBS by dialysis.

Antibody gene mutations were introduced by QuikChange II site directed mutagenesis kit (Agilent, Cat. No. 200524)

### Production of Fab from IgG

Fab fragments were produced from purified IgGs of monoclonal antibodies 2-15, 2-43 and H4 by digestion with Papain in the presence of the reducing agent 30 mM cysteine and were purified by affinity chromatography on protein A following the manufacturer’s protocols (Thermo fisher). The purity of the resultant Fabs was analyzed by SDS-PAGE, and buffer exchanged into SEC buffer (10 mM Tris pH 7.4 and 150 mM NaCl) for crystallization experiments.

### Binding Affinity Measurements by Surface Plasmon Resonance

The binding affinities of antibodies to SARS-CoV-2 spike protein were determined using surface plasmon resonance (SPR) and a BIAcore T200 instrument (GE Healthcare) at 25°C. The anti-his antibody was first immobilized onto two different flow cells of a CM5 sensorchip (BR100030, Cyvita) surface using the His Capture Kit (28995056, Cyvita) according to the manufacturer’s protocol. The His-tagged SARS-CoV-2 spike protein was then injected and captured on flow cells 2. Flow cells 1 was used as the negative control. A three-fold dilution series of antibodies with concentrations ranging from 300 nM to 1.2 nM were injected over the sensor surface for 55 seconds at a flow rate of 50 µl/minute. The dissociation was monitored for 300 seconds and the surface was regenerated with 10 mM Glycine pH 1.5 (BR100354, Cyvita). The running sample buffer is 10 mM HEPES pH 7.4, 150 mM NaCl, 3 mM EDTA, 0.05% P-20 (HBS-EP+ buffer, BR100826, Cyvita). The resulting data were fit to a 1:1 binding model using Biacore Evaluation Software and were plotted using Graphpad (Graphpad Prism 7.01, San Diego).

### Pseudovirus neutralization assays

Recombinant Indiana VSV (rVSV) expressing SARS-CoV-2 spikes were generated as previously described (Han et al., 2020, Liu et al., 2020). HEK293T cells were grown to 80% confluency before transfection with pCMV3-SARS-CoV-2-spike (kindly provided by Dr. Peihui Wang, Shandong University, China) using FuGENE 6 (Promega). Cells were cultured overnight at 37°C with 5% CO_2_. The next day, medium was removed and VSV-G pseudo-typed ΔG-luciferase (G*ΔG-luciferase, Kerafast) was used to infect the cells in DMEM at an MOI of 3 for 1 hr before washing the cells with 1X DPBS three times. DMEM supplemented with anti-VSV-G antibody (I1, mouse hybridoma supernatant from CRL-2700; ATCC) was added to the infected cells and they were cultured overnight as described above. The next day, the supernatant was harvested and clarified by centrifugation at 300g for 10 min and aliquots stored at −80°C.

Neutralization assays were performed by incubating pseudoviruses with serial dilutions of antibodies, and scored by the reduction in luciferase gene expression. In brief, Vero E6 cells were seeded in a 96-well plate at a concentration of 2 × 10^4^ cells per well. Pseudoviruses were incubated the next day with serial dilutions of the test samples in triplicate for 30 mins at 37°C. The mixture was added to cultured cells and incubated for an additional 24 hrs. The luminescence was measured by Britelite plus Reporter Gene Assay System (PerkinElmer). IC_50_ was defined as the dilution at which the relative light units were reduced by 50% compared with the virus control wells (virus + cells) after subtraction of the background in the control groups with cells only. The IC_50_ values were calculated using non-linear regression in GraphPad Prism.

### Antibody gene assignments and genetic analyses

The 158 SARS-COV-2 neutralizing antibodies were collected from ten publications. We annotated these antibodies using IgBLAST-1.16.0 with the default parameters (Ye et al., 2013). For antibodies which have cDNA sequences deposited, the V and J genes were assigned using SONAR version 2.0 (https://github.com/scharch/sonar/) with germline gene database from IMGT (Schramm et al., 2016, Lefranc et al., 2009). For each antibody, the N-addition, D gene, and P-addition regions were annotated by IMGT V-QUEST (Brochet et al., 2008). To identify somatic hypermutations, each antibody sequence was aligned to the assigned germline gene using MUSCLE v3.8.31 (Edgar, 2004). Somatic hypermutations were identified from the alignment. In addition, the analysis of single cell antibody repertoire sequencing data of SARS-CoV-2 patient 2 from (Liu et al., 2020), from which 2-15 and 2-43 were isolated, showed that 29 of the 38 unique transcripts assigned to IGLV2-14*01 share nucleotide mutations G156T and T165G. These mutations lead to amino acid mutations E50D and N53K. Both nucleotide mutations are also observed in 82 of 90 unique IGLV2-14 transcripts from patient 1 of the same study. Because these transcripts having different VJ recombination and paired with different heavy chain genes, the chances that the two convergent mutations are the results of somatic hypermutation are very low. Thus, we suspect that both donors contain a new IGVL2-14 gene allele (IGVL2-14*0X), which was deposited to European Nucleotide Archive (ENA) with project accession numbers: PRJEB31020. 2-43 and 2-15 were assigned to the IGLV2-14*0X allele.

### Cryo-EM data collection and processing

For mAb 2-43, SARS-CoV-2 spike protein at a concentration of 2 mg/ml was incubated with six-fold molar excess per spike trimer of the antibody Fab fragments for 30 minutes in 10 mM sodium acetate pH 5.5, 150 mM NaCl, and 0.005% n-dodecyl-β-D-maltoside (DDM). 2µL of the sample was incubated on C-flat 1.2/1.3 carbon grids for 30 seconds and vitrified using a Leica EM GP. Data were collected on a Titan Krios electron microscope operating at 300 kV, equipped with a Gatan K3 direct electron detector and energy filter, using the Leginon software package (Suloway et al., 2005). A total electron fluence of 51.69 e-/Å2 was fractionated over 40 frames, with a total exposure time of 2.0 seconds. A magnification of 81,000x resulted in a pixel size of 1.058 Å, and a defocus range of −0.4 to −3.5 µm was used.

For mAb 2-15, SARS-CoV-2 spike protein at a concentration of 1 mg/ml was incubated with a molar ratio of 1:1 Fab fragments to spike trimer for 30 minutes in 10 mM sodium acetate pH 5.5, 150 mM NaCl, and 0.005% % DDM. 2µL of the sample was incubated on C-flat 1.2/1.3 carbon grids for 30 seconds and vitrified using a Leica EM GP. Data were collected on a Titan Krios electron microscope operating at 300 kV, equipped with a Gatan K3 direct electron detector and energy filter, using the Leginon software package (Suloway et al., 2005). A total electron fluence of 52.40 e-/Å2 was fractionated over 60 frames, with a total exposure time of 3.0 seconds. A magnification of 81,000x resulted in a pixel size of 1.070 Å, and a defocus range of −0.8 to −3.4 µm was used.

For mAb H4, SARS-CoV-2 spike protein at a concentration of 1 mg/ml was incubated with eight-fold molar excess per spike trimer of the antibody Fab fragments for 30 minutes in 10 mM sodium acetate pH 5.5, 150 mM NaCl, and 0.005% % DDM. 2µL of the sample was incubated on C-flat 1.2/1.3 carbon grids for 30 seconds and vitrified using a Leica EM GP. Data were collected on a Titan Krios electron microscope operating at 300 kV, equipped with a Gatan K3 direct electron detector and energy filter, using the Leginon software package. A total electron fluence of 42.00 e-/Å2 was fractionated over 40 frames, with a total exposure time of 3.0 seconds. A magnification of 81,000x resulted in a pixel size of 1.070 Å, and a defocus range of −0.5 to −3.5 µm was used.

All processing was done using cryoSPARC v2.15.0 (Punjani et al., 2017). Raw movies were aligned and dose-weighted using patch motion correction, and the CTF was estimated using patch CTF estimation. A small subset of approximately 200 micrographs were picked using blob picker, followed by 2D classification and manual curation of particle picks, and used to train a Topaz neural network. This network was then used to pick particles from the remaining micrographs, which were extracted with a box size of 384 pixels.

For the mAb 2-43 dataset, 2D classification followed by ab initio modelling and 3D heterogeneous refinement revealed 61,434 particles with three Fabs bound, one to each RBD. A reconstruction of these particles with imposed C3 symmetry resulted in a 3.78 Å map, as determined by gold standard Fourier shell correlation (FSC). Symmetry expansion followed by masked local refinement was used to obtain a 3.81 Å map of the Fab and RBD. The remainder of the S trimer was subjected to local refinement to obtain a 3.61 Å map. These two separate local refinements were aligned and combined using the vop maximum function in UCSF Chimera (Pettersen et al., 2004). This was repeated for the half maps, which were used along with the refinement mask from the global non-uniform refinement to calculate the 3D FSC (ref) and obtain an estimated resolution of 3.60 Å. All maps have been submitted to the EMDB with the ID EMD-#####.

For the mAb 2-15 dataset, 2D classification followed by ab initio modelling and 3D heterogeneous refinement revealed 16,590 particles with one Fab bound to an RBD in the ‘down’ conformation and 21,456 particles with one Fab bound to an RBD in the ‘up’ conformation. The particles with Fab bound to RBD down were refined using Non-uniform refinement and C1 symmetry to a global resolution of 5.73 Å as determined by gold standard FSC. The RBD and Fab were masked and subjected to local refinement to obtain a map at 6.21 Å. The remainder of the trimer was also refined locally to 5.63 Å. A consensus map was obtained as described for mAb 2-43, with a resolution of 5.87 Å. All maps have been submitted to the EMDB with the ID EMD-#####.

For the mAb H4 dataset, 2D classification followed by ab initio modelling and 3D heterogeneous refinement revealed 102,290 particles with one Fab bound to an RBD in the ‘up’ conformation. No classes with Fab bound to ‘down’ RBD were identified. 3D Variability Analysis was used to visualize the significant conformational heterogeneity of the ‘up’ RBD. Using the first and last frames of the reaction coordinate as reference maps, representing the extremes of the orientations adopted by the RBD, heterogeneous refinement was repeated to separate the Fab-bound spikes into more homogeneous classes. 56,080 particles adopted a more stable conformation and were refined to 4.78 Å using homogeneous refinement and C1 symmetry. Like the previously described datasets, the Fab and RBD were refined locally to 5.03 Å, with the remainder of the S trimer being refined to 4.32 Å. The final consensus map was estimated to have a resolution of 5.07 Å. All maps have been submitted to the EMDB with the ID EMD-#####.

### Cryo-EM model building

An initial homology model of the 2-43 Fab was built using Schrödinger Release 2020-2:BioLuminate (Zhu et al., 2014) and of H4 using ABodyBuilder (Leem et al., 2016). For mAb 2-15, the crystal structure determined here (PDB ID ####) was used as a starting model for the Fab variable domain and the associated RBD. For all models, the S trimer was modeled using the coordinates from PDB ID 6XEY. These models were docked into the consensus map using Chimera. The model was then fitted interactively using ISOLDE 1.0.1 (Croll, 2018) and COOT 0.8.9.2 (Emsley and Cowtan, 2004) and using real space refinement in Phenix 1.18 (Adams et al., 2004). For Fab 2-43, in cases were side chains were not visible in the experimental data, they were truncated to alanine except for residues very close to the RBD:Fab interface. Both H4 and 2-15 were built as poly-alanine models due to the low resolution of the experimental data. Validation was performed using Molprobity (Davis et al., 2004) and EMRinger (Barad et al., 2015). Models were submitted to the PDB with the following IDs: mAb 2-43 is 7L56, mAb H4 is 7L58, and 2-15 is 7L57. Figures were prepared using UCSF ChimeraX (Goddard et al., 2018).

### X-ray data collection, structure solution, model building and refinement

For determination of the complex of with RBD, the two proteins were mixed at 1:1 molar ratio and incubated at 4.0°C for 60 min. RBD:2-15 complex was then isolated by gel filtration on Superdex-200 (GE Healthcare). Fractions containing complexes were pooled and concentrated to 6.5 mg/ml in SEC buffer. Screening for initial crystallization conditions was carried out in 96-well sitting drop plates using the vapour-diffusion method with a Mosquito crystallization robot (TTP LabTech) using various commercially available crystallization screens. Diffracting crystals were obtained from 0.1 M Hepes pH 7.5 and 70% MPD. Crystals were directly frozen and X-ray diffraction data was collected to 3.18 Å resolution at 100 K from a single flash-cooled crystal on beamline ID-C at the Advance Photon Source (APS) at Argonne National Laboratory. Diffraction data were processed with XDS (Kabsch, 2010) and scaled using AIMLESS (Evans, 2006) from the CCP4 software suite (Collaborative Computational Project, 1994). Molecular replacement was performed with Phaser (McCoy et al., 2007), with previously reported RBD structure (PDB 7BZ5) and Fab (PDB code, 4HK0) as search models. Manual rebuilding of the structure using COOT (Emsley et al., 2010) was alternated with refinement using Phenix refine (Afonine et al., 2012) and PDB-REDO (Joosten et al., 2014). The Molprobity server was used for structure validation and PyMOL (version 2.1, Schrödinger, LLC) for structure visualization (Chen et al., 2010). A summary of the X-ray data collection and refinement statistics are shown in Table S2. PISA was used to identify paratope-epitope interfaces and to calculate buried surface area (Krissinel and Henrick, 2007). Hydrogen bonds were identified with a cutoff of 3.8Å. 2-15 model was submitted to the PDB with the following IDs: 7L5B.

### Calculation of angle of approach

To measure the RBD approaching angle of antibodies, we first identify shared epitope residues among the five members of the VH1-2 antibody class. PyMOL was used to determine the centre of mass of the shared epitope residues. We then determined the center of mass for heavy (the centre of mass of the conserved Cys at 22 and 92) and light chains (the centre of mass of the conserved Cys at 23 and 88). The heavy and light chain approaching angle was determined by linking the chain centre of mass to the centre of mass of the shared epitope. PyMOL was used to make structure figures. For antibody B38, the epitope center of mass was determined using the epitope residues of the antibody.

## QUANTIFICATION AND STATISTICAL ANALYSIS

The statistical analyses for the pseudovirus neutralization assessments were performed using GraphPad Prism. The SPR data were fitted using Biacore Evaluation Software. Cryo-EM data were processed and analyzed using CryoSparc and Relion. Cryo-EM structural statistics were analyzed with Phenix and Molprobity. Statistical details of experiments are described in Method Details or figure legends.

