## Supplementary figures and images for "Modular basis for potent SARS-CoV-2 neutralization by a prevalent VH1-2-derived antibody class"

### Figure S1

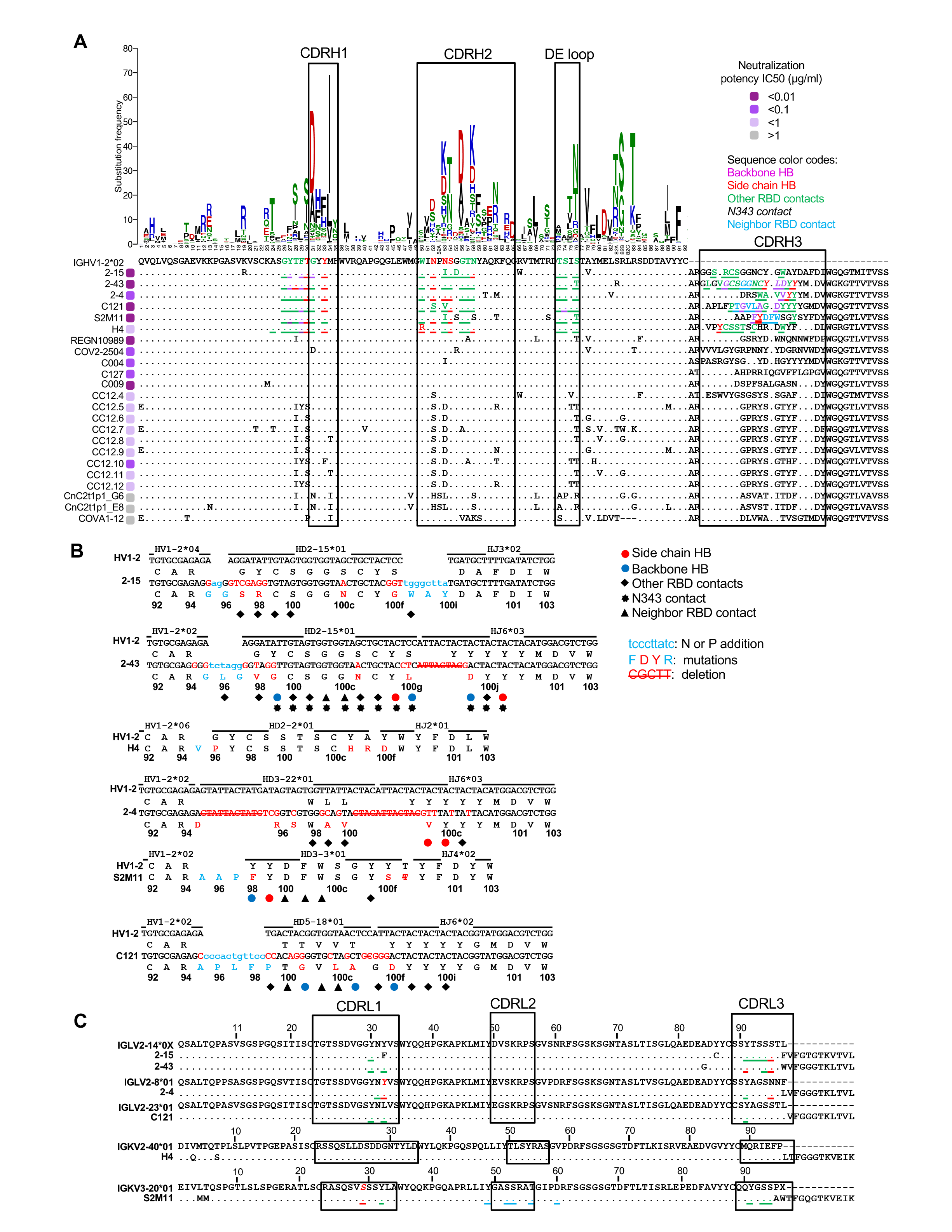

### Figure S1

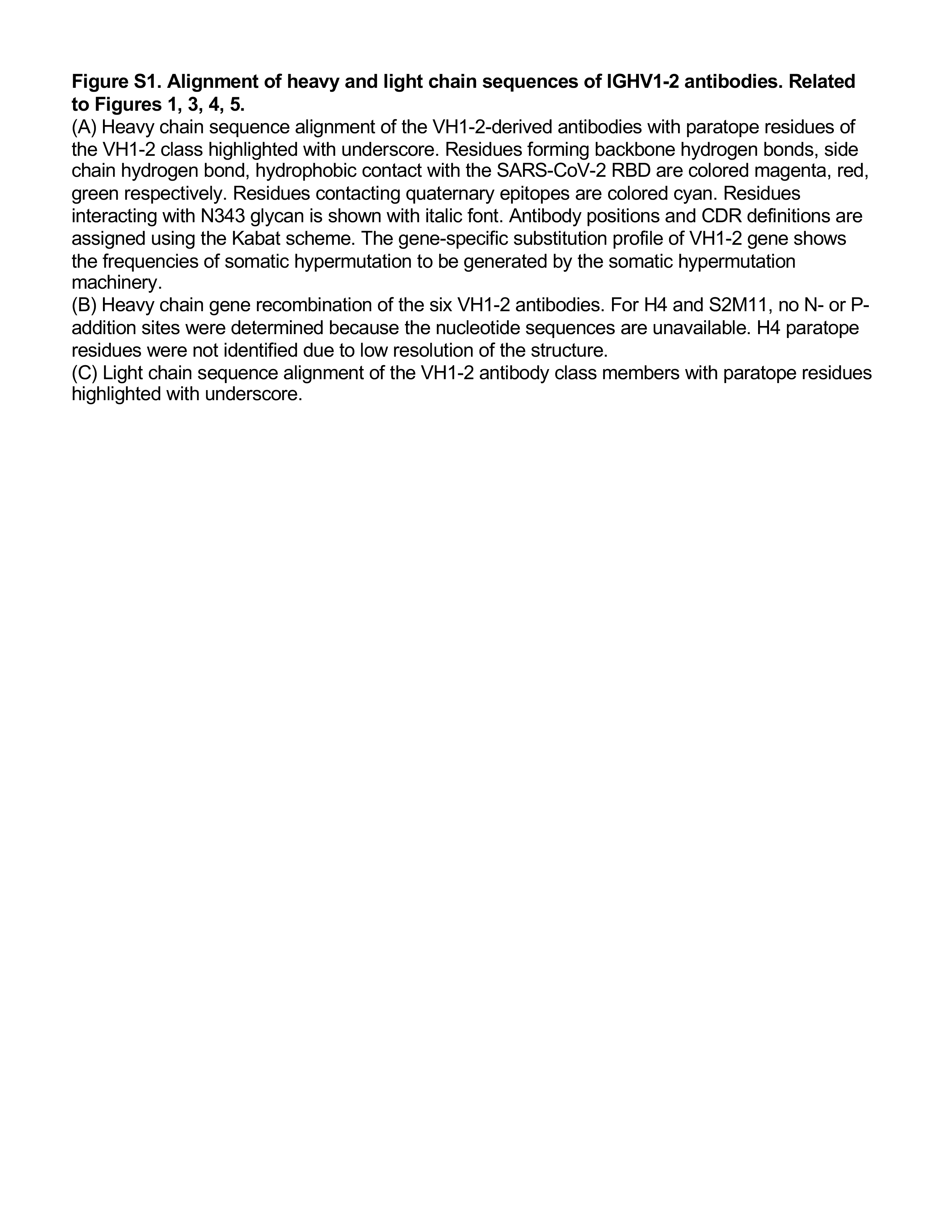

### Figure S2

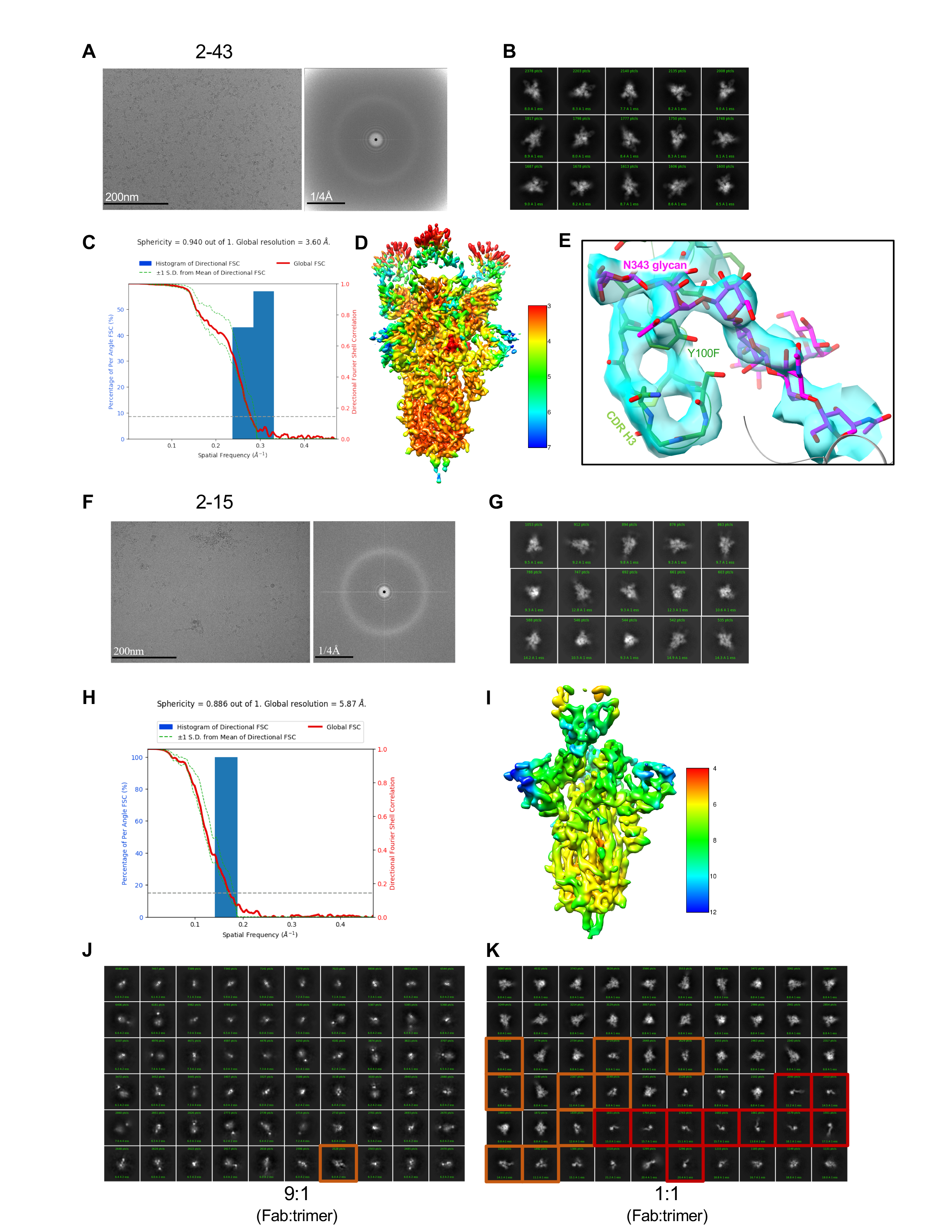

### Figure S2

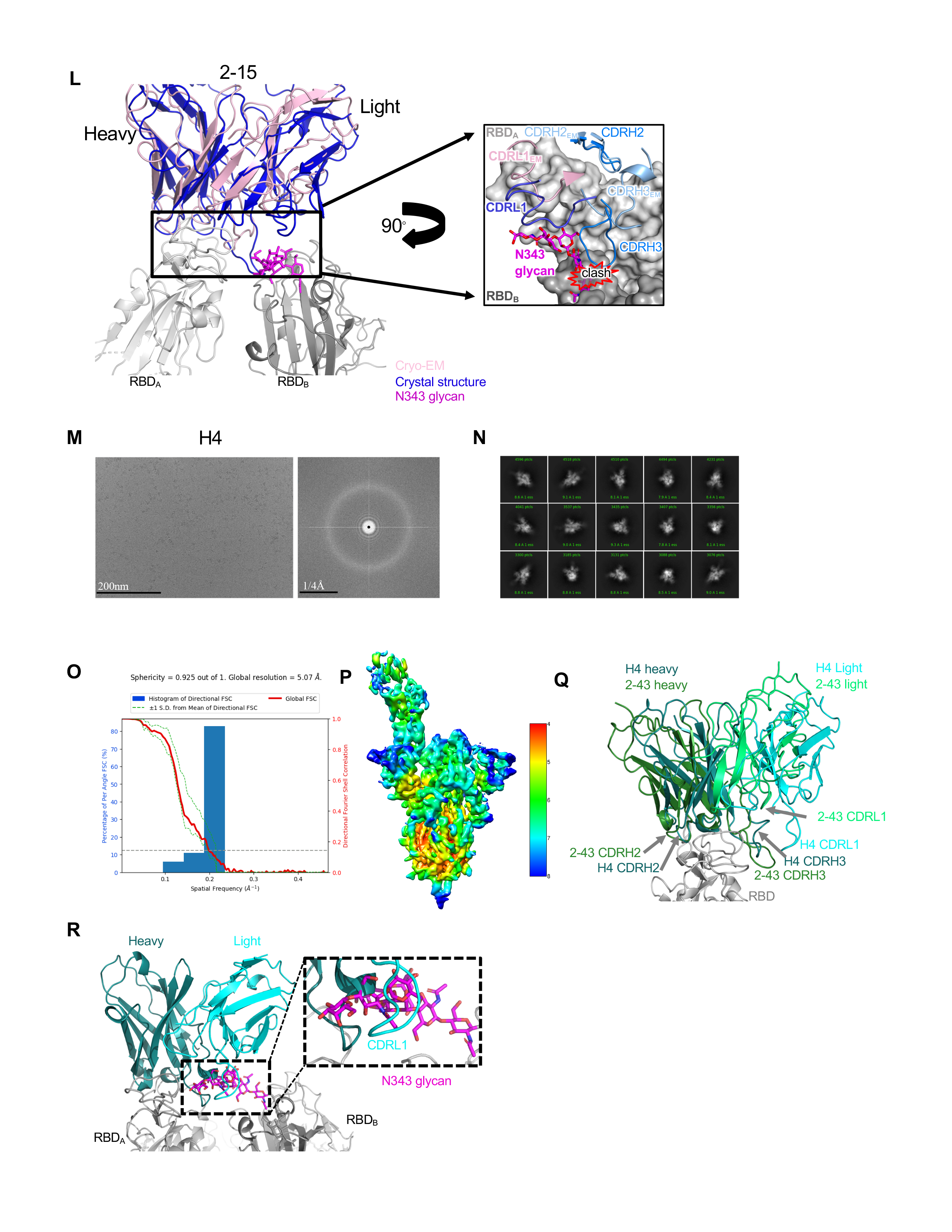

### Figure S2

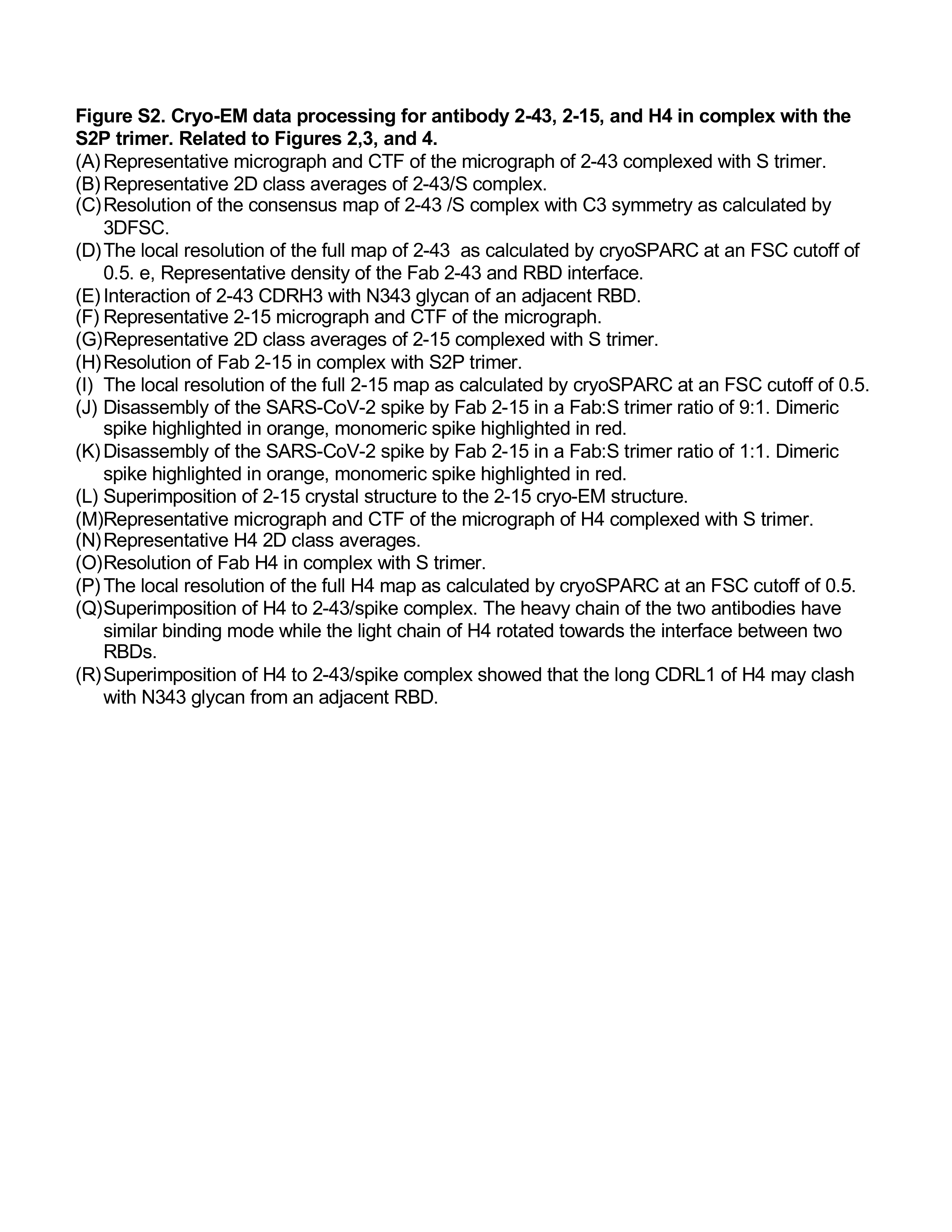

### Figure S3

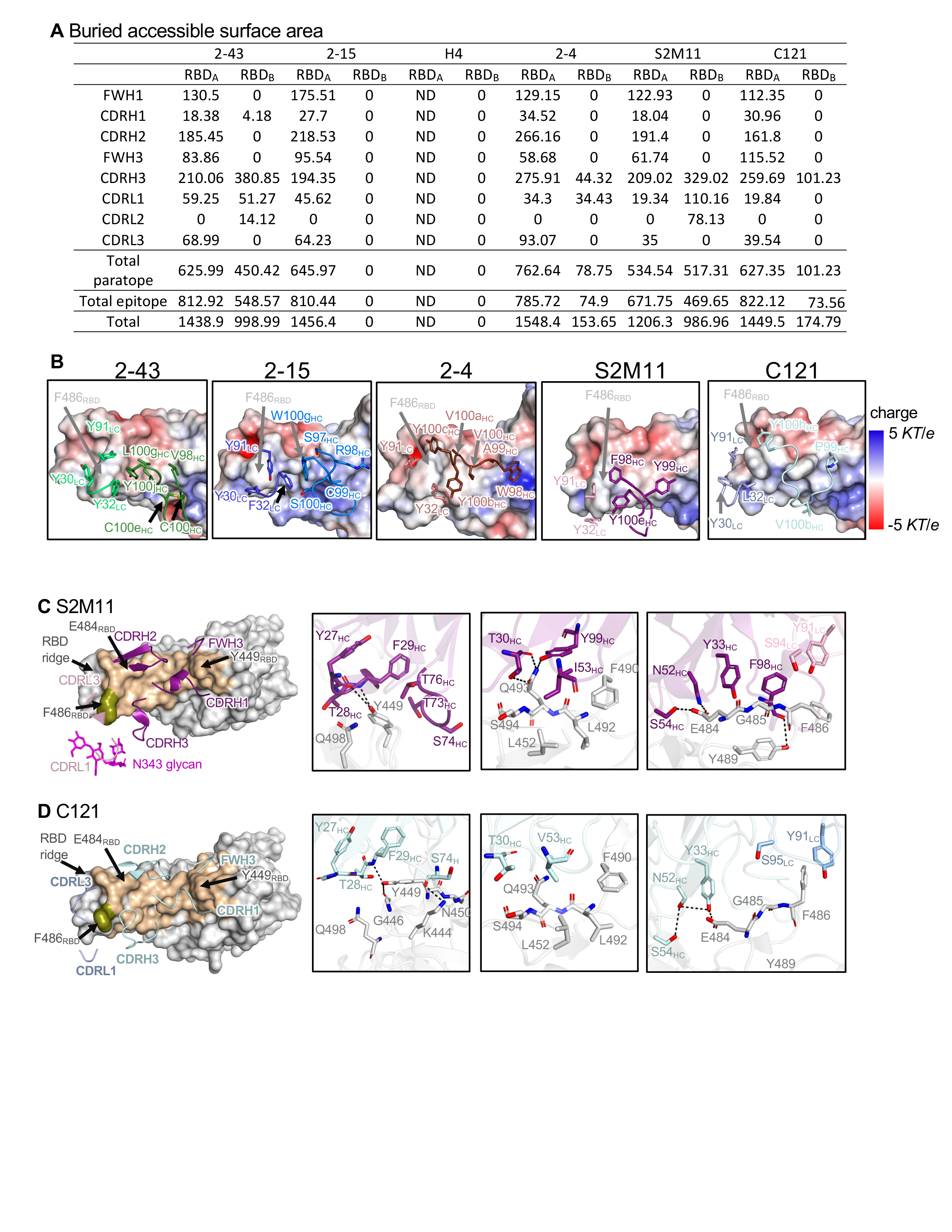

### Figure S3

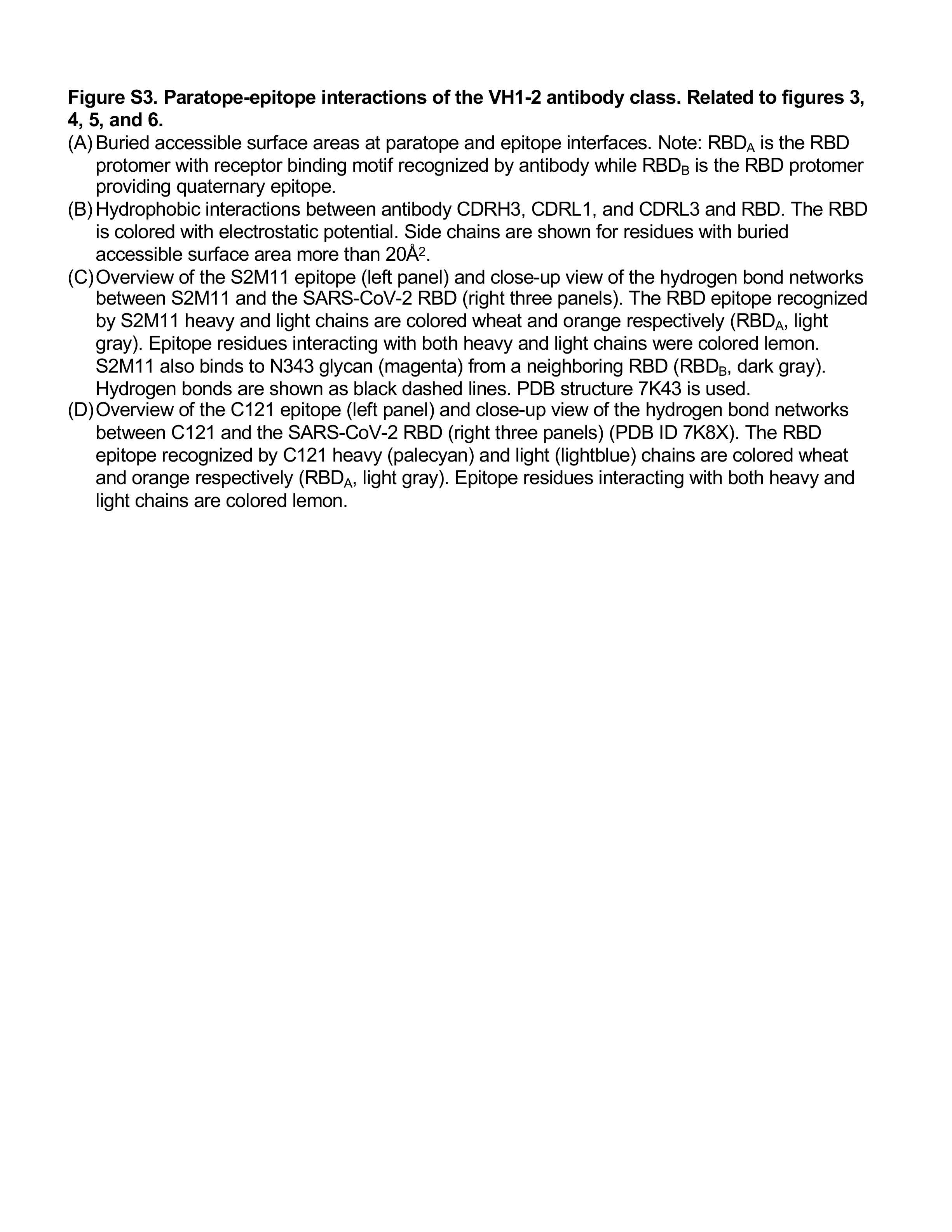

### Figure S4

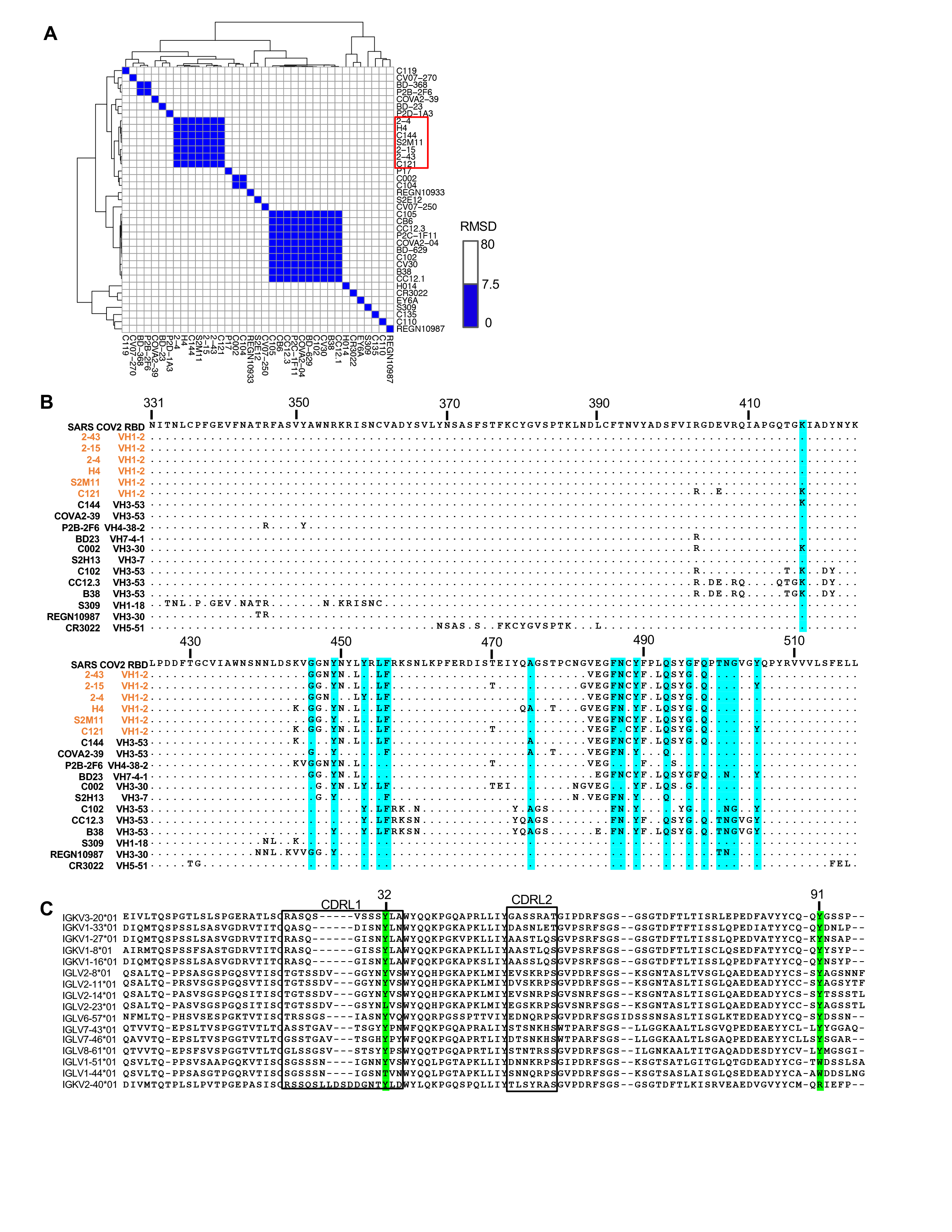

### Figure S4

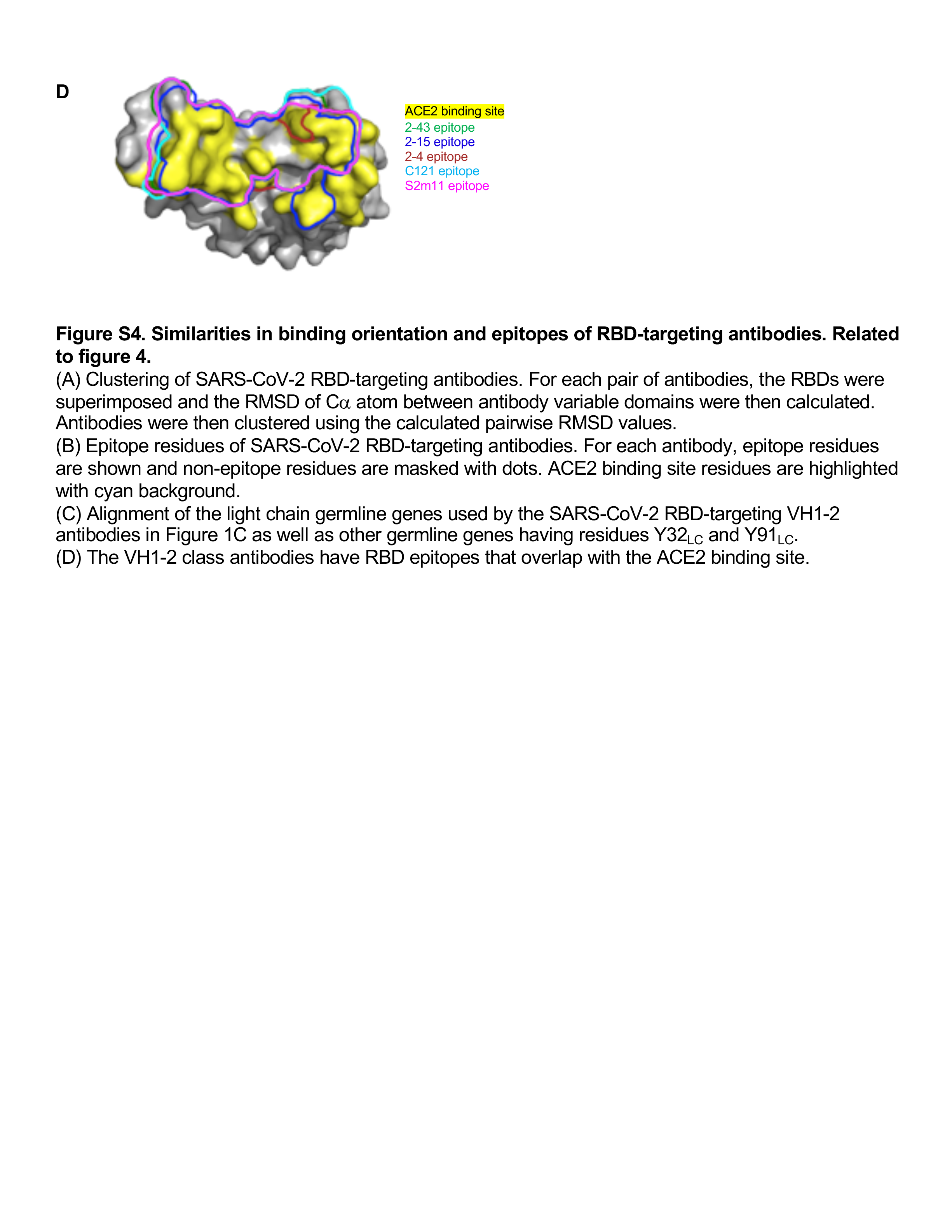

### Figure S5

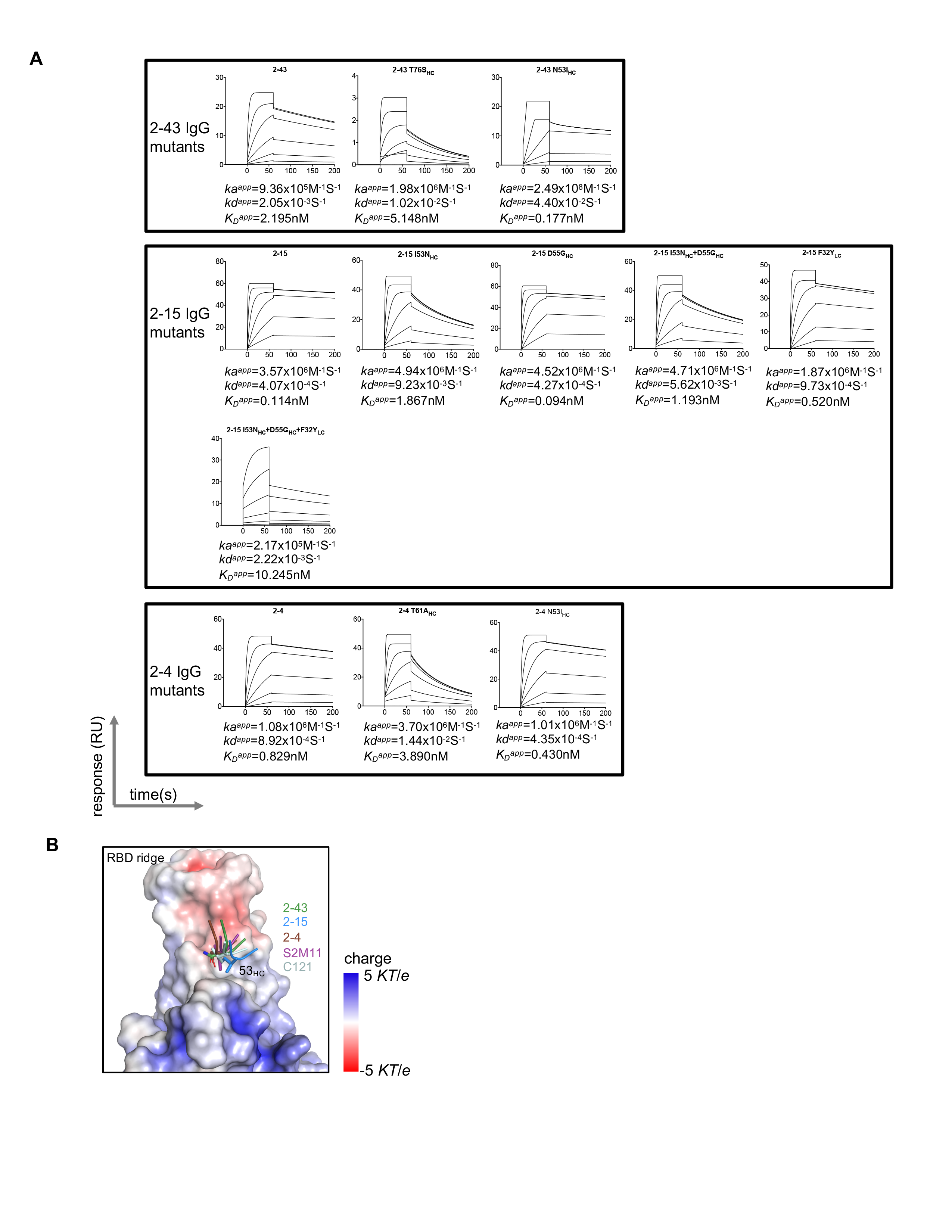

### Figure S5

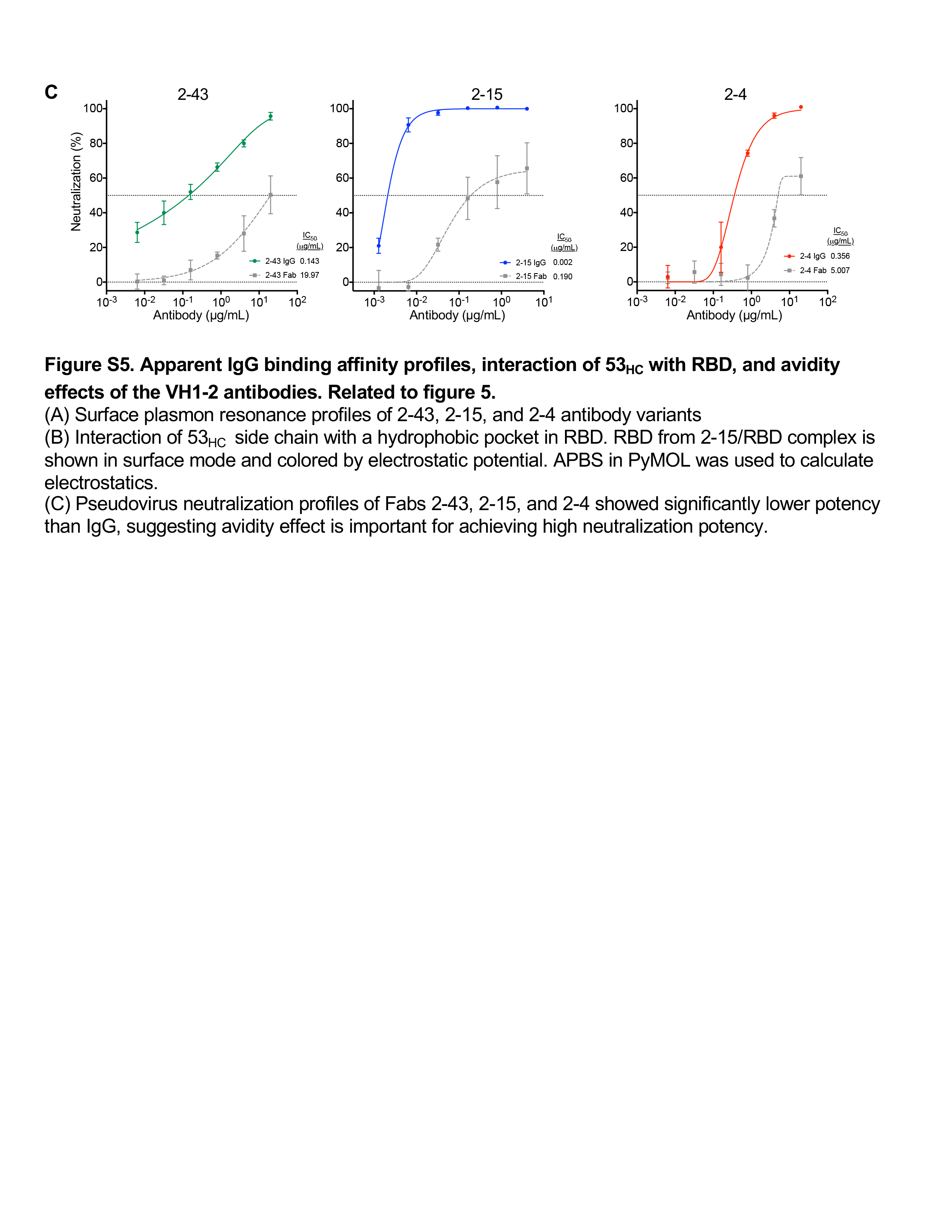

### Figure S6

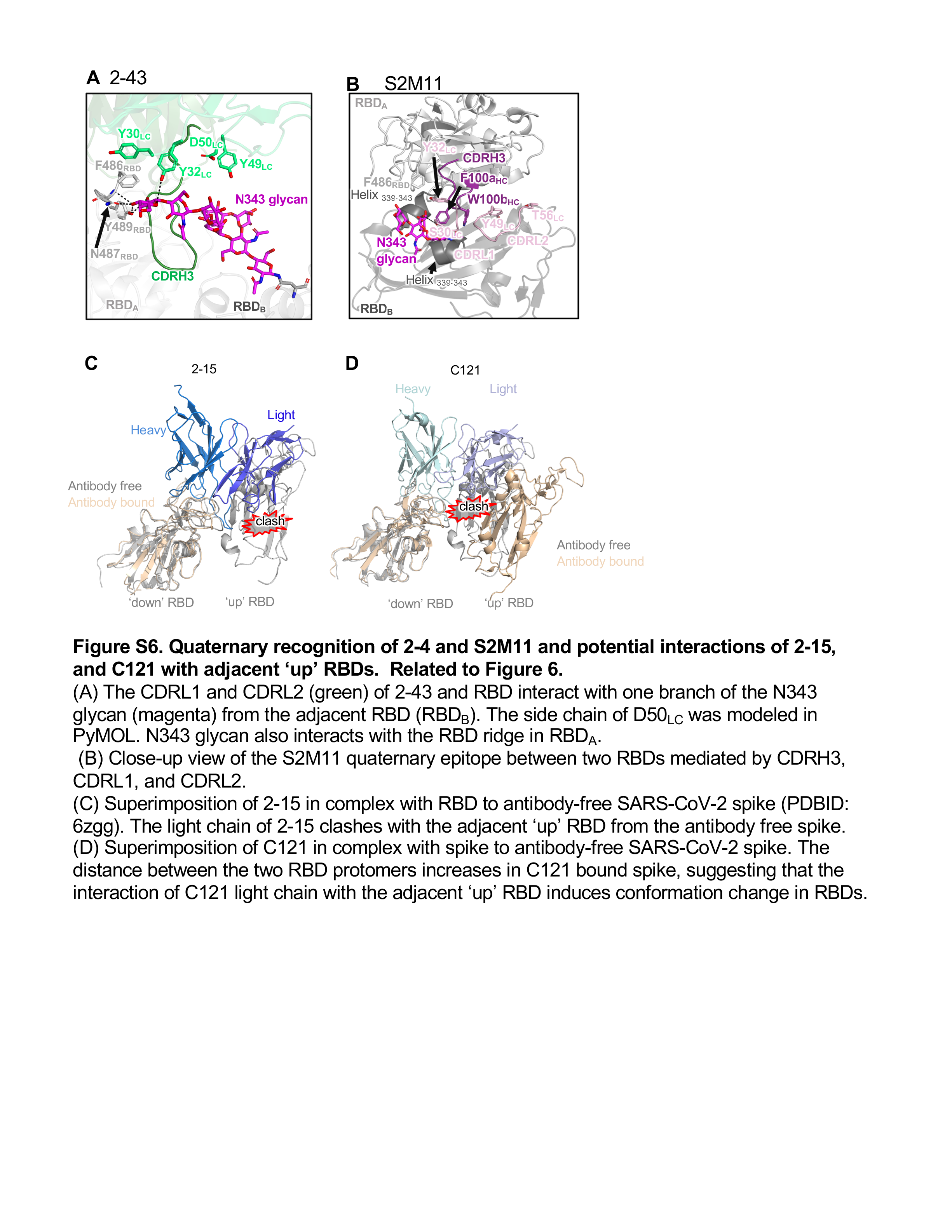

### Table S1

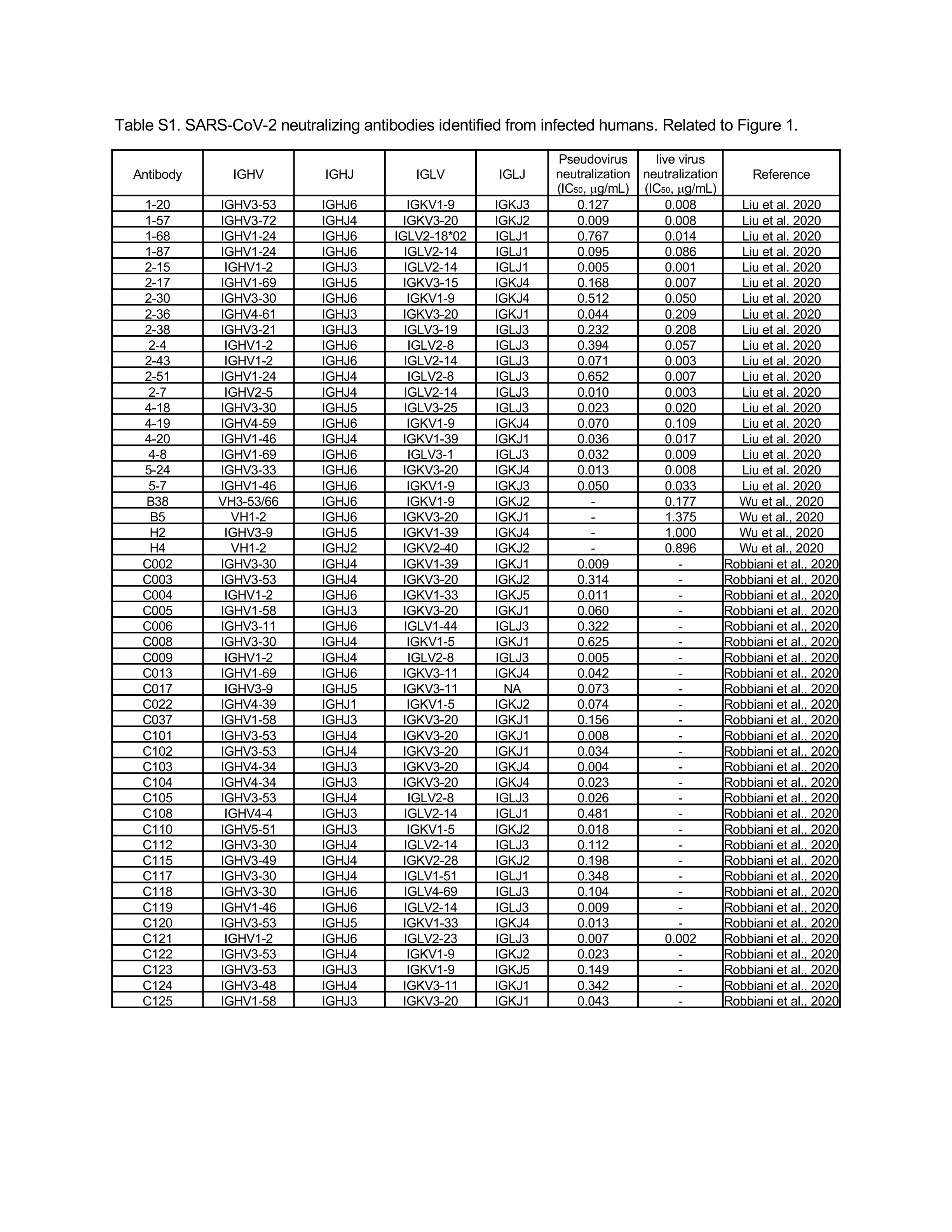

### Table S1

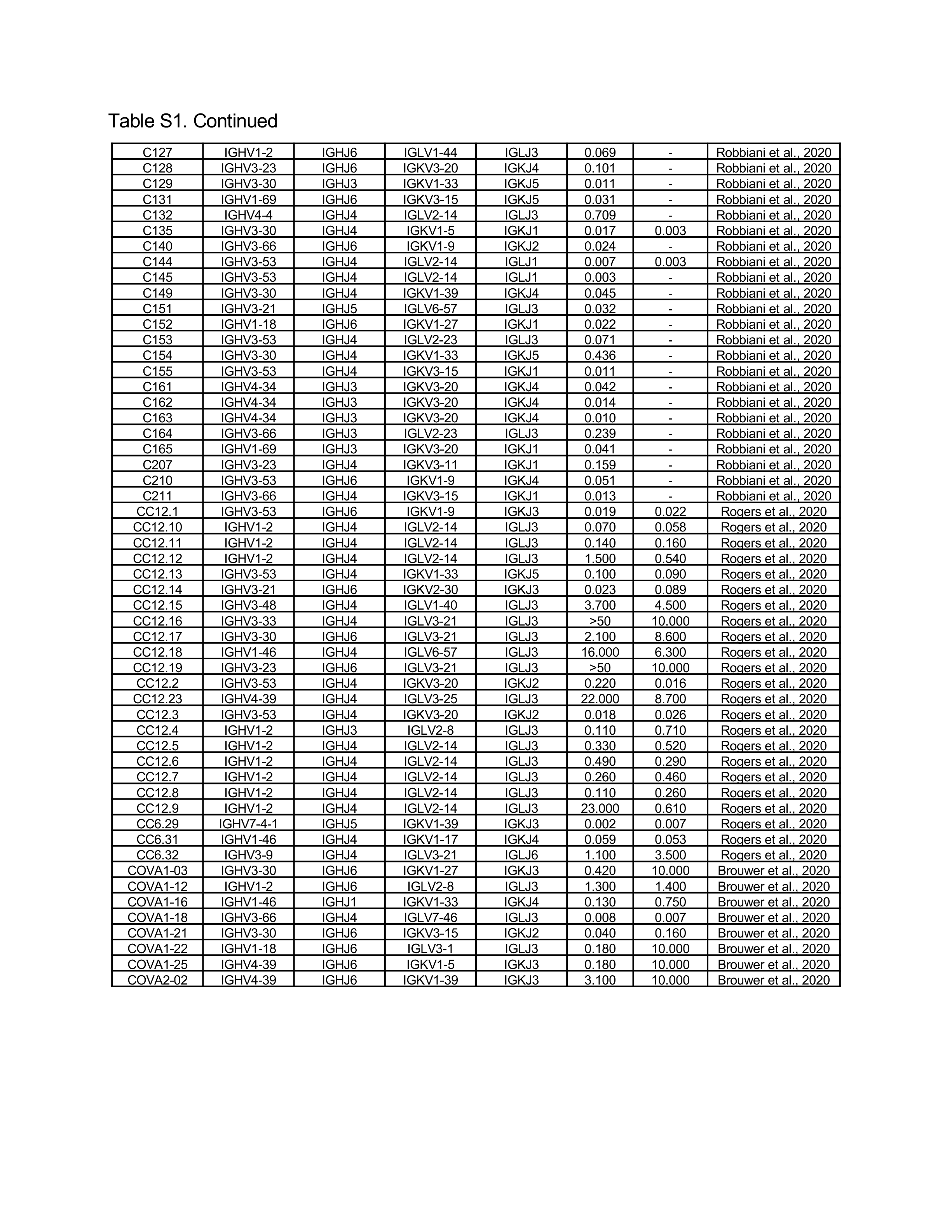

### Table S1

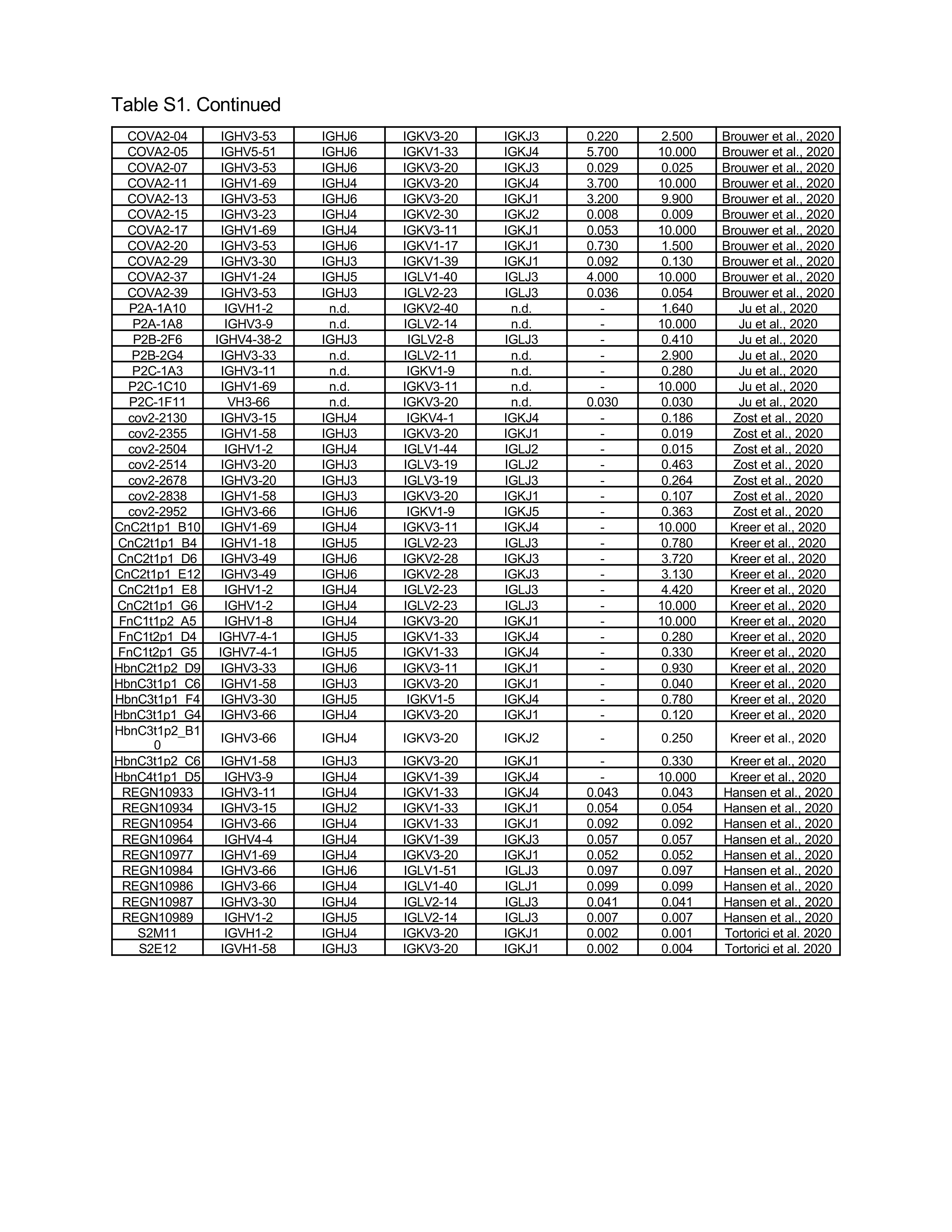

### Table S2

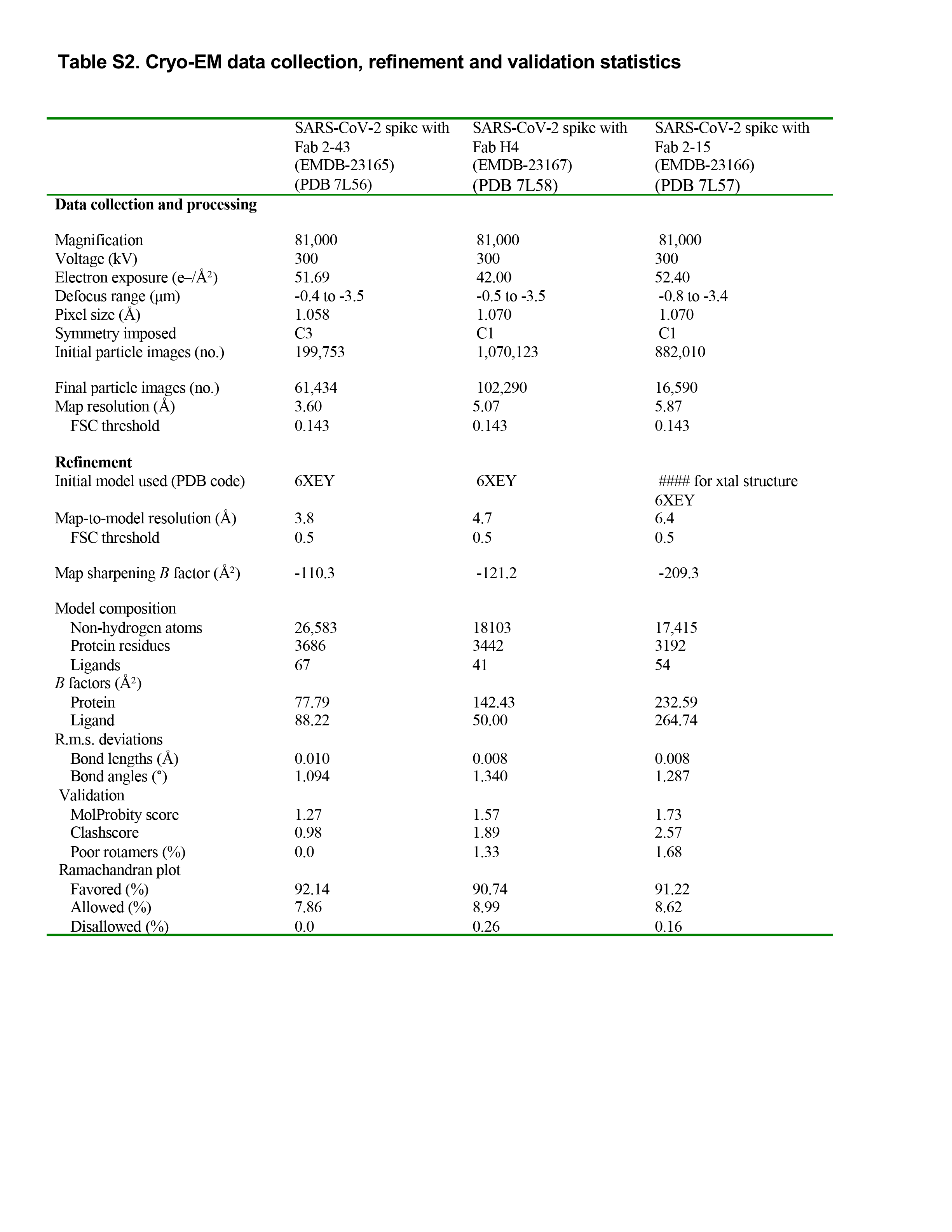

### Table S3

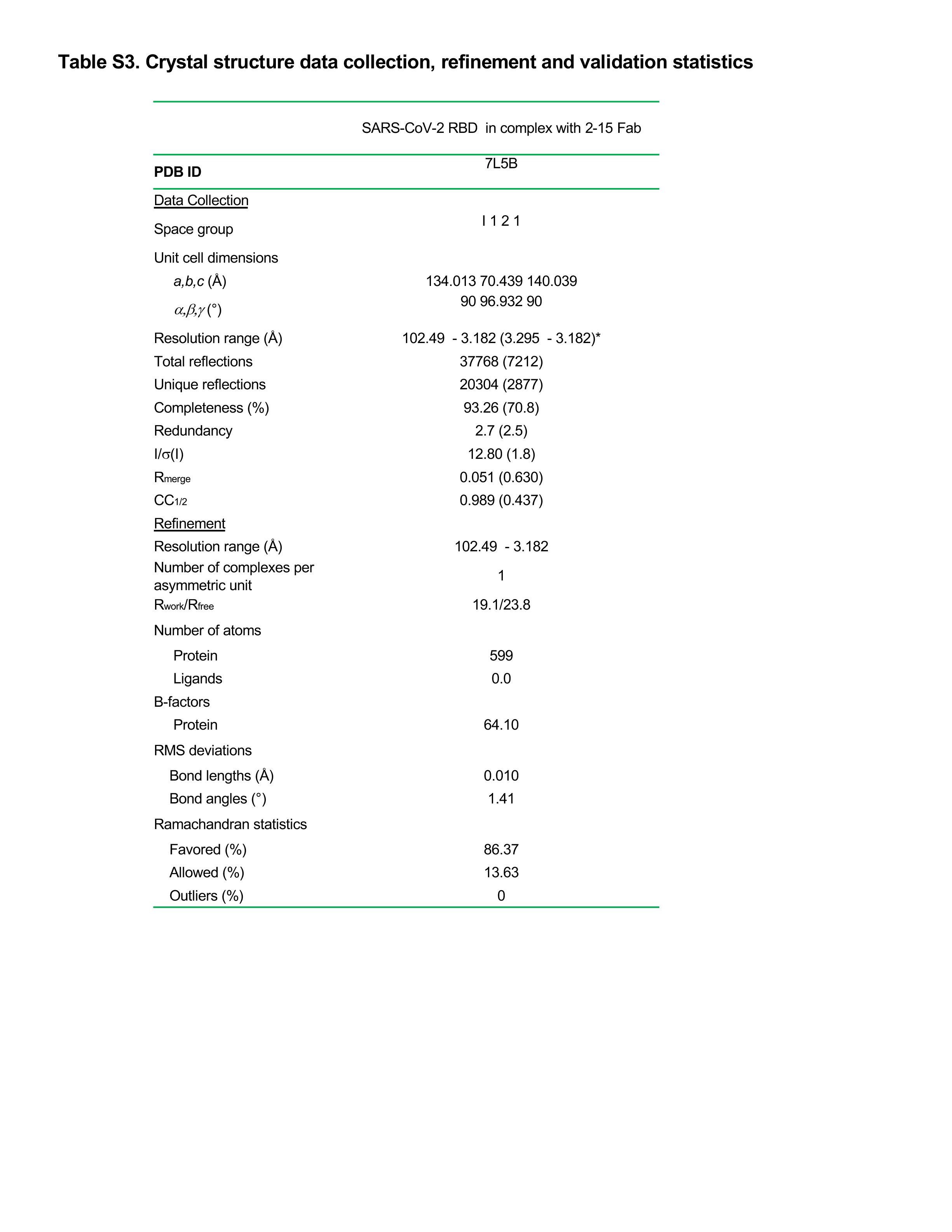
